# Drought responses reshape growth, reproduction and seed traits in *Camelina sativa*

**DOI:** 10.64898/2026.11.25.769118

**Authors:** Miguel Benavides-Acevedo, Annika Grieß-Osowski, Jenny Köpplin, Santana Saam, Björn Usadel, Carolina Lobato

## Abstract

*Camelina sativa* is an emerging oilseed crop for low-input and climate-resilient agriculture, valued for its agronomic versatility and tolerance to abiotic stress. However, how drought affects its seeds, and whether these effects carry over to the next generation, remains poorly understood. We grew six *Camelina* accessions at 100% (control) and 20% (drought) soil water content and followed their responses from vegetative growth to seed mucilage content and germination. Drought reduced stomatal conductance and increased leaf temperature in all accessions, whereas its effect on shoot growth changed over time and differed among accessions. Drought-stressed plants developed a more compact shoot, with shorter secondary stems, inflorescences and internodes and 43% higher secondary stem density. Seed yield per plant fell by 47%, but seeds were 38% heavier and 28% larger in all accessions, consistent with plants prioritizing the provisioning of individual seeds over seed number. These larger seeds had a 27% smaller mucilage area relative to their size, suggesting that drought shifts maternal allocation away from the seed coat, which could affect seedling establishment. Drought also shifted the monosaccharide composition of seed mucilage, mainly in the non-adherent fraction, where rhamnose increased and galactose decreased in only one accession (CAM230), whereas in the adherent fraction it lowered the proportion of galacturonic acid. Despite their larger size, progeny of drought-stressed plants germinated less under salt (NaCl) and osmotic (PEG) stress, and under PEG (−0.6 MPa) germination fell by up to 26 percentage points in half of the accessions. These results reflect differences among accessions in how shoot growth responds to drought over time and a cost of maternal drought for the progeny. Both should be considered when selecting *Camelina* for drought-prone, low-input systems.

## Introduction

Drought events are projected to become more frequent and severe in some regions as the climate warms (IPCC, 2021), and agriculture absorbs almost 80% of their direct economic impacts worldwide (Food and Agriculture Organization of the United Nations, 2023). Drought alters plant metabolism, growth and reproduction (Basu et al., 2016; Chaves et al., 2003), and the environment of the mother plant can also shape the phenotype of its offspring (Donohue, 2009; Herman and Sultan, 2011). The conditions during seed production, for example, affect progeny dormancy and germination through maternal abscisic acid, the seed coat and dormancy regulators such as DOG1 (Penfield and MacGregor, 2017). Transgenerational effects are adaptive when the parental environment induces offspring phenotypes that perform better in that same environment (Herman and Sultan, 2011). In the halophyte *Suaeda salsa*, for example, progeny of mother plants grown under high salinity showed better seedling emergence, growth and seed production (Guo et al., 2020). Maternal drought also affects the progeny by decreasing seed dormancy (Nicotra et al., 2010), and in *Brassica napus* changes in seed provisioning and storage metabolites have been proposed to underlie its transgenerational effects (Hatzig et al., 2018).

Seed mucilage is one route by which the mother plant can affect its progeny. Many Brassicaceae are myxospermous, so when their seeds imbibe, polysaccharides stored in the epidermal cells of the seed coat are extruded as mucilage (Haughn and Western, 2012; North et al., 2014). Unlike the embryo, the seed coat develops from the maternal integuments (Arsovski et al., 2010; Haughn and Western, 2012), so mucilage is produced entirely by maternal tissue. Mucilage can be separated into an adherent layer, which remains attached to the seed, and a non-adherent layer, which detaches readily (Miart et al., 2019; Voiniciuc et al., 2016). In *Arabidopsis*, the adherent layer is held to the seed by xylan side chains on rhamnogalacturonan I, which bind the mucilage to cellulose (Ralet et al., 2016). Mucilage has been proposed to support germination under salt and osmotic stress, to retain water and to anchor seeds or affect their dispersal, although these effects vary among species (Bhatt et al., 2016; Engelbrecht et al., 2014; Tsai et al., 2021; Yang et al., 2021). In *Plantago* species, mucilage quantity increases toward hotter and drier sites of origin, which suggests a role in adaptation to water deficit (Teixeira et al., 2020). In *Arabidopsis*, by contrast, natural variants with altered mucilage release produce seeds that float, but an adaptive advantage of this trait has not been demonstrated (Saez-Aguayo et al., 2014).

*Camelina sativa* (*Camelina*) is an emerging oilseed crop of the Brassicaceae that can be grown on marginal land and has potential for drought-resilient agriculture; its oil-rich seeds are mainly used for biofuel (Zanetti et al., 2024). As a close relative of *Arabidopsis*, *Camelina* also extrudes seed mucilage with adherent and non-adherent layers, but its mucilage is richer in arabinose and galactose and varies widely among accessions (Perera et al., 2022). Despite relatively low genetic diversity (Brock et al., 2024), *Camelina* shows substantial phenotypic plasticity and genotype × environment interactions in response to abiotic stress, including drought (Alberghini et al., 2025; Silvestre et al., 2026). Drought research in Camelina has focused on how the mother plant performs, from its physiology and growth to seed yield and quality (Alberghini et al., 2025; Silvestre et al., 2026; Vayssières et al., 2026; Waraich et al., 2017), and seeds of different sizes have been tested for germination under osmotic stress (Čanak et al., 2020). Whether maternal drought alters the amount and composition of seed mucilage, and how such changes relate to seed size and to the germination of the next generation, has not been examined.

We therefore tested two hypotheses: (H1) maternal drought affects mucilage production, mucilage composition and seed traits; and (H2) mucilage production covaries with seed traits and with the germination of the progeny under stress. We also asked whether these responses are consistent among accessions. To address these questions, we grew six Camelina accessions under control and drought conditions from before bolting until harvest and followed the mother plants from leaf physiology and shoot growth to seed yield. We then traced the drought effect into their progeny seeds, where we measured how much mucilage was extruded relative to seed size, how its composition differed between the adherent and non-adherent layers, and how well the seeds germinated under salt and osmotic stress.

## Materials and methods

### Seed collection and plant-growing conditions

*Camelina sativa* (L.) Crantz accessions (CAM22, CAM70, CAM131, CAM190, CAM214 and CAM230) were obtained from the IPK seedbank (Gatersleben, Germany). Seeds were stratified for 72 hours at 4 °C, and then placed in a germination tray with 193-grade germination paper (Sartorius) for 14 days inside a climate cabinet (Poly Klima L2-TDL+) with a constant temperature of 22 °C, 60% humidity and 16/8 light cycle. Grown seedlings were transplanted into 2 L pots filled with 517 g (± 13 g) of standard horticultural peat-based substrate (D400) and coconut coir (CocoPor), and supplemented with the controlled-release fertilizer Osmocote 5 (3-4M; 16-8-12 + 2.2 MgO + trace elements) at a rate of 3 g L^−1^ of substrate. A total of 36 plants (6 plants per accession) were grown under greenhouse conditions at the Heinrich-Heine University Düsseldorf (Germany) between May and August 2024 with a 16/8 light cycle (**Figure S1A**). The temperature was 22 °C, and supplementary lamps gave a photosynthetically active radiation of about 200 µmol m^−2^ s^−1^ at the base of the pot and 230 µmol m^−2^ s^−1^ at the top of the plant, in addition to natural daylight. Plants were randomized and watered to field capacity three times a week until the end of 4 weeks after sowing (WAS), before bolting. At that point, three plants of each accession were watered to 100% of soil-water content (SWC), while the remaining plants were watered to 20% SWC until the end of the life cycle. SWC was determined by weighing substrate-filled pots at oven-dry (0%) and field capacity (100%), where the weight difference was used to calculate target water contents.

### Plant growth and physiology

Plant growth (projected shoot area) was monitored from the start of the drought treatment (4 WAS) to 9 WAS using image-based phenotyping (**Figure S1B**). Plants were imaged weekly in a controlled PhotoBox using an Olympus E-M1 Mark III camera (12–45 mm 1:4 PRO lens). For each plant four lateral images were acquired, rotating the plant by 90° between images. Image processing was performed using Color_Segmentation_Tool_V4pub (Mueller-Linow, 2022) and Fiji (Schindelin et al., 2012). Color-based segmentation was applied to isolate plant tissue, using HSV thresholds of Hue = 32–57, Saturation = 41–255, and Value = 36–255 for fully green plants. For plants exhibiting chlorosis or senescence, thresholds were adjusted to Hue = 12–57, Saturation = 41–255, and Value = 36–255 to ensure accurate segmentation. Subsequent image analysis was automated in Fiji using a custom macro (Benavides-Acevedo, 2025a), including conversion to 8-bit grayscale, automatic thresholding to generate binary masks and particle analysis to identify plant objects (minimum size ≥0.30 mm²), and extraction of area measurements. The mean projected shoot area was calculated from the four images per plant at each time point. Leaf temperature and stomatal conductance to water vapor (gsw) were measured at 7 WAS using a Li-600 porometer. Measurements were performed on two leaves per plant spanning lower and upper canopy positions (approx. the 10th to the 20th leaf). For each leaf, temperature and gsw were recorded at basal, middle, and apical regions.

### Plant architecture and reproduction

Architectural traits were measured after 13 WAS, following complete senescence of inflorescences. The basal main stem diameter was measured in the first centimeters of the main stem immediately above the soil surface, whereas the mid main stem diameter was measured at approximately 50% of the total main stem height. The main stem diameter ratio (basal diameter/mid-stem diameter), was calculated to describe variation in stem tapering, with higher values indicating a greater reduction in stem diameter from the base toward the middle portion of the stem. Internode distance was measured between consecutive secondary stems and secondary stem length recorded (**Figure S1B**).

Main stem length was measured from the point where the stem emerged from the substrate to the uppermost branching point of the main stem, which corresponded to the node where the main stem inflorescence originated. Secondary stem density was calculated as the number of secondary stems per cm of main stem length. Reproductive traits and yield were quantified at 13 WAS to characterize inflorescence development and allocation patterns. The secondary stem main inflorescence length (SSMIL) was measured from the node where the stem inflorescence originated (i.e., the uppermost branching point of the stem) to the apex of the inflorescence, to quantify variation in reproductive structure development across the plant (**Figure S1B**). Seeds were collected from each plant and weighed to obtain seed yield per plant (g) (**Figure S1C**).

### Seed traits

Thousand-seed weight (g) was determined by weighing approximately 1,000 counted seeds per plant and seed number per plant was calculated as seed yield divided by the mass of one seed (**Figure S1C**). Seeds were imaged using an Olympus camera under uniform backlighting provided by an ultra-slim RGB CCT LED panel (29.5 × 29.5 cm; 3000–6500 K; 1800 lumens) adapted as a negatoscope. Seed size was quantified using Fiji (Schindelin et al., 2012) and a custom ImageJ macro (Benavides-Acevedo, 2025a). Image processing included conversion to 8-bit grayscale, automated thresholding (Default method), application of a fixed intensity range (0-179), binary mask generation, and particle analysis (minimum size ≥0.30 mm²) to isolate individual seeds and extract area measurements. Progeny seeds were stored in paper bags in the dark at room temperature and germinated under controlled conditions using distilled water (control), NaCl (122.3 and 244.5 mM; −0.6 and −1.2 MPa), and polyethylene glycol (PEG-8000) (−0.4, −0.6, −0.8 and −1.0 MPa) twelve months after harvesting. For each maternal plant and condition, 30 seeds were germinated in 90 mm Petri dishes containing 193-grade germination paper (Sartorius) moistened with 2 mL of the corresponding solution, and 1 mL of fresh solution was added daily. Seeds were incubated in a growth chamber (Poly Klima L2-TDL+) at 22 °C, 75% relative humidity, and continuous light (24-hour photoperiod). Germination, defined as radicle emergence through the seed coat, was counted daily for 3 days.

### Seed mucilage

For mucilage area, progeny seeds were sampled two months after harvest from three stems per plant, including the main inflorescence of the main stem (M), and the main inflorescences of one basal (B), and one top (T) secondary stem counted from the base of the main stem. Within each stem, seeds were taken from three different capsules. Seeds were imbibed following the MuSeeQ protocol (Miart et al., 2018), with three seeds per sample in distilled water for 24 h at room temperature. Images were acquired under identical backlighting conditions and processed using Color_Segmentation_Tool_V4pub (Mueller-Linow, 2022) and Fiji (Schindelin et al., 2012). Seed size was segmented in *LabCH* color space, while total seed plus mucilage area was segmented using HSV thresholding. Image analysis was automated using a custom Fiji macro, including ROI cropping, image alignment (3° rotation), conversion to 8-bit grayscale, and thresholding to generate binary masks. Particle analysis (minimum size ≥0.30 mm²) was used to identify objects and extract standardized measurements (Benavides-Acevedo, 2025b). Mucilage area per seed (MAS) was calculated as the difference between the total area and the seed size, from which mucilage-to-seed size ratio was derived (**Figure S1C**). Monosaccharide composition of seed mucilage was determined using the protocol described by Voiniciuc (Voiniciuc, 2016), modified for *Camelina* seeds. Briefly, non-adherent mucilage (NAM) was recovered from the supernatant after centrifugation. The remaining pellets were washed, and adherent mucilage (AM) was extracted by sonication (Branson SFX 250) with the following settings: 10 s ON / 1 s OFF; 45 s total; 10% amplitude. Samples were dried at 45 °C under nitrogen stream and hydrolyzed with trifluoroacetic acid (TFA) at 120 °C for 60 min. Following evaporation and isopropanol washes, monosaccharides were resuspended and analyzed using ion chromatography on a Thermo Dionex ICS 6000 system with NaOH gradient elution. The analysis targeted arabinose (Ara), galactose (Gal), galacturonic acid (GalA), glucose (Glc), mannose (Man), rhamnose (Rha) and xylose (Xyl). Seed samples (7–9 mg) were incubated with ribose (15 µg/mL) as an internal standard. Seed samples from the basal, middle and apical capsules of the main inflorescence were extracted separately. Monosaccharide composition was expressed as molar percentage of the total of the seven analyzed monosaccharides in each sample and averaged per plant and mucilage fraction (157 samples from 35 plants for adherent and 34 plants for non-adherent mucilage).

### Statistical analysis

All analyses were performed in R v4.5.3 (R Core Team, 2026), with the maternal plant as the experimental unit and accession, treatment and their interaction as fixed effects unless stated otherwise. Traits with one value per plant (number of secondary stems, main stem length, secondary stem density, basal-to-mid stem diameter ratio, seed yield, seed number and seed weight) were analyzed with linear models (LMs), and traits measured on several stems, seeds or leaves per plant with linear mixed-effects models (LMMs) in lme4 v2.0.6 (Bates et al., 2015), with plant as a random intercept. Stomatal conductance, which varied much less under drought, was analyzed with nlme v3.1.170 (Pinheiro and Bates, 2000) with separate residual and between-plant variances per treatment. Projected shoot area was analyzed with glmmTMB v1.1.14 (Brooks et al., 2017), with accession × treatment × week (A × T × W) as fixed effects and a first-order autoregressive correlation between consecutive weeks. Stem-class differences in MAS and the MAS : seed size ratio were tested with accession × treatment × stem class (A × T × S) as fixed effects and stem within plant as an additional random intercept. Secondary stem density and the MAS : seed size ratio were also analyzed by analysis of covariance (ANCOVA) of the number of secondary stems on main stem length (LM) and of log MAS on log seed size (LMM), with centred covariates and a covariate × treatment interaction to check that slopes did not differ between treatments. These mixed models were fitted by restricted maximum likelihood (REML). Fixed effects were tested with Type III tests and sum-to-zero contrasts: F tests in car v3.1.5 (Fox and Weisberg, 2019) for LMs, F tests with Satterthwaite degrees of freedom in lmerTest v3.2.1 (Kuznetsova et al., 2017) for LMMs, and Wald χ² tests for shoot area. Residuals were checked with DHARMa v0.5.0 (Hartig, 2026).

Germination (seeds germinated out of 30 per plant) was analyzed with binomial generalized linear mixed models (GLMMs) in blme v1.0.8 (Chung et al., 2013), with accession, treatment, condition and their two-way interactions as fixed effects and plant and observation-level random intercepts, the latter to account for overdispersion. Accession-specific drought effects were tested in each condition with a likelihood-ratio test of the accession × treatment (A × T) interaction. Effects are reported as odds ratios (OR). Associations with germination were tested in each stress condition by adding plant means of seed size, seed weight or MAS (with seed size as covariate) to binomial GLMMs with accession and treatment (lme4; likelihood-ratio tests, including MAS × treatment), and among accessions by Spearman correlations between drought changes. Monosaccharides were related to germination in the same two ways, and to seed size and weight with LMs (plants) and Spearman correlations (accessions). P values were FDR-adjusted across traits, monosaccharides, fractions and conditions.

Monosaccharide molar percentages were analyzed on plant means with one LM per monosaccharide and fraction. Fraction differences and treatment × fraction interactions were tested with LMs on the per-plant difference between fractions. Treatment P values were adjusted across the 14 monosaccharide × fraction combinations with the false discovery rate (FDR; Benjamini and Hochberg, 1995). Overall composition was analyzed on centred log-ratio (clr) transformed data (Aitchison, 1986) by principal component analysis and by permutational multivariate analysis of variance (PERMANOVA; Anderson, 2001) on Euclidean distances with sequential terms and 9,999 permutations, using adonis2 in vegan v2.7.5 (Oksanen et al., 2026); homogeneity of dispersion was checked with betadisper.

Treatment was compared within accessions, and within weeks, stem classes or fractions where relevant, with estimated marginal means in emmeans v2.0.4 (Lenth and Piaskowski, 2026) and Holm adjustment (Holm, 1979) across accessions or, for stem classes, across all 18 accession × stem class contrasts. Data were processed and plotted with tidyverse v2.0.0 (Wickham et al., 2019) and patchwork v1.3.2 (Pedersen, 2025). Test statistics for all models are given in Table S1.

## Results

### Drought disrupts growth dynamics in an accession-dependent manner and activates physiological stress responses

To characterize the responses of *Camelina* to drought, we followed the projected shoot area of six accessions from the start of the drought treatment (4 WAS) to 9 WAS and measured leaf temperature and stomatal conductance (**Figure 1**). The drought response of shoot area changed over time and differed between accessions (A × T × W, P < 0.001), so treatment effects were tested within each accession and week. At the start of the treatment (4 WAS), plants assigned to drought were on average slightly larger than control plants (+68 cm², 95% CI 18-119 cm², P = 0.008), although the difference was significant only in CAM230 (+178 cm², P = 0.029). After one week of drought (5 WAS), drought-treated plants were smaller on average (−112 cm², 95% CI −162 to −61 cm², P < 0.001), but no single accession differed significantly (P ≥ 0.068). The reduction peaked at 6 WAS, when shoot area was smaller under drought in all six accessions (by 222.5-450.5 cm²; P < 0.001 for each accession). Thereafter, the drought effect declined as shoot area of control plants decreased after its peak, and accessions diverged: CAM22, CAM214 and CAM230 remained significantly smaller under drought until 9 WAS (by 183.3-258.4 cm²; P ≤ 0.015), whereas CAM70 and CAM190 no longer differed from control plants from 7 WAS onward (P ≥ 0.051), and CAM131 from 8 WAS onward (P ≥ 0.085; **Figure 1A**). Drought also altered leaf gas exchange. Leaf temperature was 1.28 °C higher under drought (95% CI 1.11-1.44 °C; P < 0.001; **Figure 1B**), and stomatal conductance fell from 0.41 to 0.02 mol m^−2^ s^−1^ (difference −0.39, 95% CI −0.48 to −0.30 mol m^−2^ s^−1^; P < 0.001; **Figure 1C**). Neither response differed significantly between accessions (A × T, P = 0.071 for leaf temperature and P = 0.36 for stomatal conductance), and both were significant in every accession (**Figure S2**).

**Figure 1.**
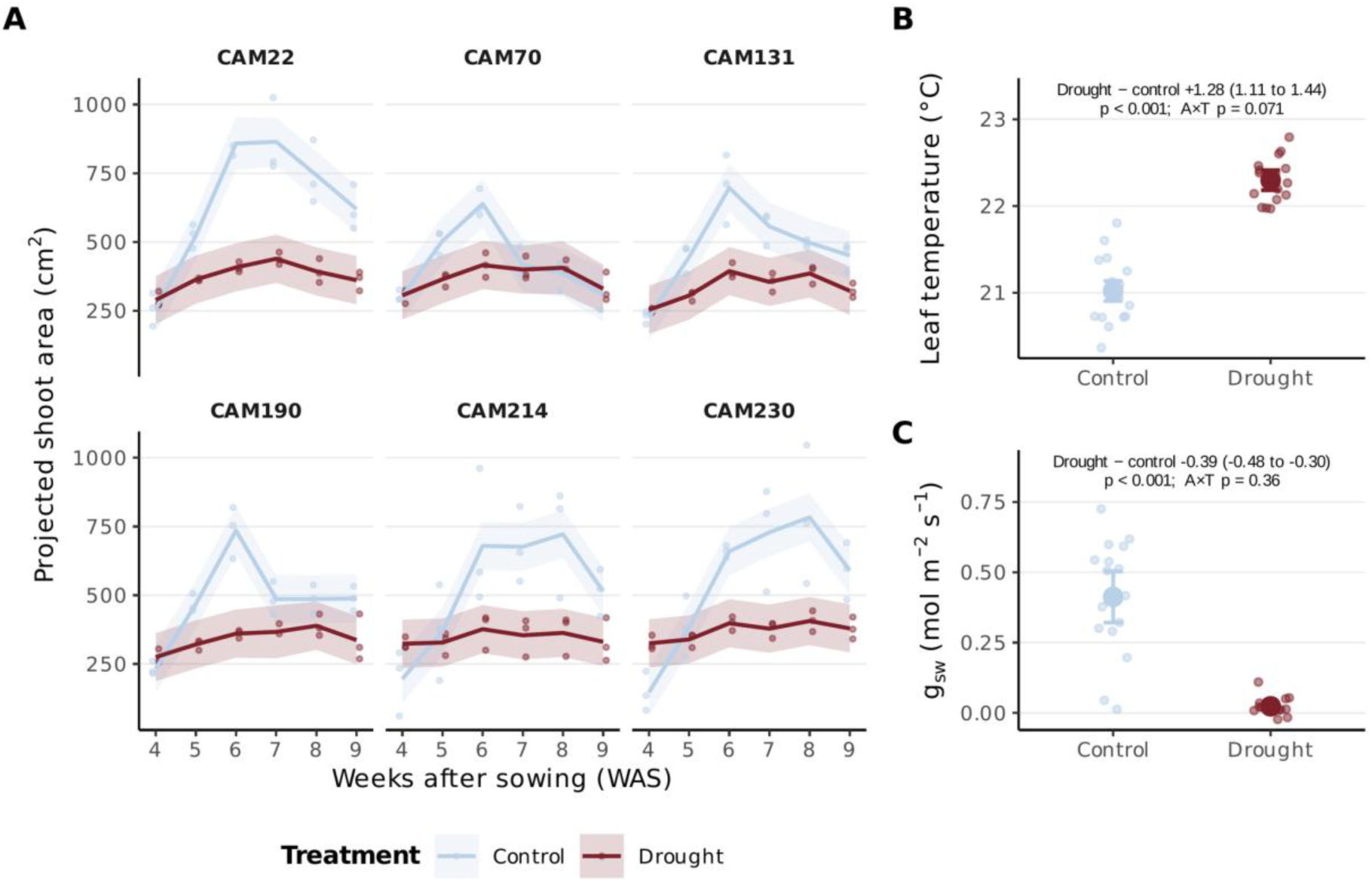
Shoot growth, leaf temperature and stomatal conductance of *Camelina* accessions under control and drought conditions. **(A)** Projected shoot area per accession over 6 weeks (4-9 WAS) under control and drought (glmmTMB with AR(1) correlation over weeks). Lines and bands are model means with 95 % CI and dots are individual plants. **(B)** Leaf temperature and **(C)** stomatal conductance (gsw) averaged over accessions (accession × treatment (A × T), P = 0.071 and 0.36). Subtitles give the drought - control difference with its 95 % CI, and dots represent plant means.

### Drought reshapes maternal plant architecture and shifts reproduction toward fewer, larger seeds

To assess how drought affects the maternal plant and the seeds it produces, we analyzed stem architecture, seed yield and seed traits of six *Camelina* accessions grown under control and drought conditions (**Figure 2**). Our results showed no evidence that accessions respond differently to drought, as the A × T interaction was not significant for any of these traits (P = 0.088-0.97). Effects for each accession are shown in **Figure S3**. Secondary stem length decreased by 52% (95% CI 42-60%; P < 0.001), SSMIL by 56% (51-61%; P < 0.001) and internode distance by 28% (23-33%; P < 0.001) under drought. Consistent with the shorter internodes, secondary stems were more densely spaced along the main stem, with secondary stem density increasing by 43% (33-53%; P < 0.001). This reflected a 27% shorter main stem (23-30%; P < 0.001) rather than more secondary stems, whose number increased only by 4.5% (1.1-8.0%; P = 0.011) and did not change significantly in any accession (**Figure S3D**). With main stem length as a covariate, drought had no detectable effect on the number of secondary stems (+1%, P = 0.85; **Table S2**). The ratio of basal to mid-stem diameter was not affected by drought (+2%, −6 to +11%; P = 0.64). Seed yield per plant decreased by 47% under drought (36-57%; P < 0.001) and seed number per plant by 61% (50-70%; P < 0.001). In contrast, the progeny seeds were heavier and larger: seed weight increased by 38% (29-47%; P < 0.001) and seed size by 28% (23-33%; P < 0.001). Drought-stressed maternal plants therefore produced fewer seeds, but each seed was larger and heavier.

**Figure 2.**
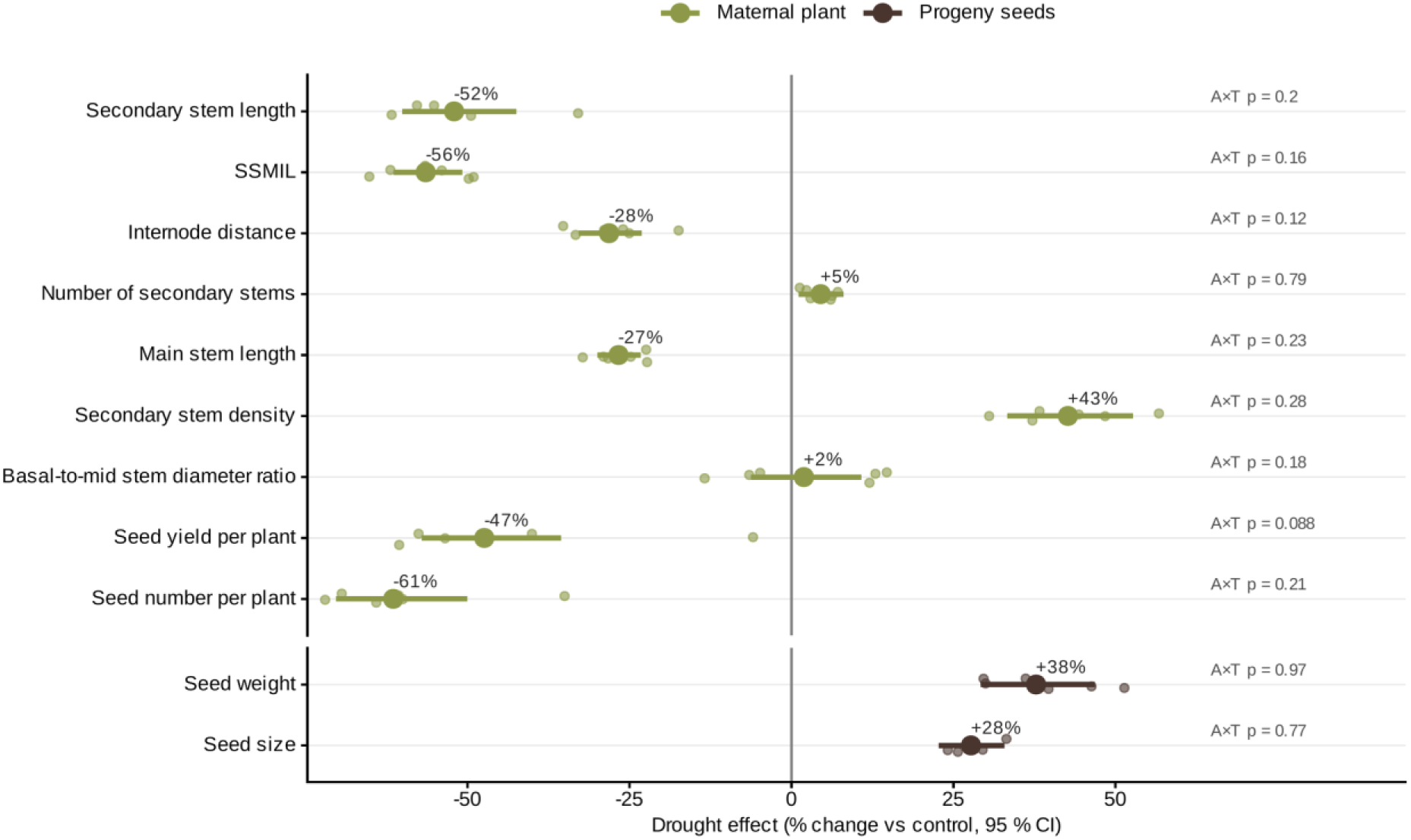
Effect of drought on maternal plant architecture, seed yield and progeny seed traits. Drought effects are shown as % change relative to the control plants of that same accession (95 % CI, delta method on the log ratio of marginal means) for traits whose accession × treatment (A × T) interaction is not significant. Large points are effects averaged over accessions, colored by trait group (maternal plant in green and progeny seeds in brown), while small points in the same color represent single accessions. SSMIL, secondary stem main inflorescence length.

### Larger seeds from drought-stressed plants carry proportionally less mucilage

Because drought produced larger seeds (**Figure 2**), we tested whether seed mucilage changed accordingly. Averaged over accessions, mucilage area per seed (MAS) did not differ between treatments (−7%, 95% CI −18 to +5%; P = 0.23), and the A × T interaction was not significant (P = 0.081; **Figure 3A**). No single accession differed significantly after Holm adjustment, although MAS tended to decrease in CAM230 and to increase in CAM131 (**Figure 3A**). With larger seeds and unchanged MAS, the MAS : seed size ratio decreased by 27% under drought (95% CI 19-35%; P < 0.001; **Figure 3B**). This reduction did not differ significantly between accessions (A × T, P = 0.10), ranging from 9% in CAM131 to 51% in CAM230, and was significant individually in CAM22 (−39%, P = 0.002) and CAM230 (−51%, P = 0.008; **Figure 3B**). We also analyzed log MAS with log seed size as a covariate (**Table S2**). Mucilage area increased more than proportionally with seed size (slope 2.24, 95% CI 2.16-2.32), meaning that at equal seed size drought seeds had 47% less mucilage area than control seeds (39-54%; P < 0.001). This reduction did not differ between accessions (A × T, P = 0.23) and was significant in each of them (37-60%, Holm P ≤ 0.012). To assess whether stem position modulates these responses, MAS and the MAS : seed size ratio were analyzed in seeds from a basal (B) and a top (T) secondary stem and from the main stem main inflorescence (M) of each plant (**Figure 3C-D**). The drought response did not depend on stem class: neither the treatment × stem class interaction (P = 0.63 for MAS and 0.21 for the ratio) nor the A × T × S interaction (P = 0.83 and 0.82) was significant. Accordingly, MAS was unaffected by drought in all three stem classes (P ≥ 0.094; **Figure 3C**), whereas the ratio was lower under drought in each of them (by 2.2-3.0; P < 0.001; **Figure 3D**). Differences between stem classes, however, varied among accessions irrespective of treatment (accession × stem class, P = 0.048 for MAS; P = 0.008 for the ratio). For example, MAS was highest in basal stems in CAM230 but lowest in basal stems in CAM22 and CAM70, and CAM214 had the lowest MAS in top stems (**Figure S4A**).

**Figure 3.**
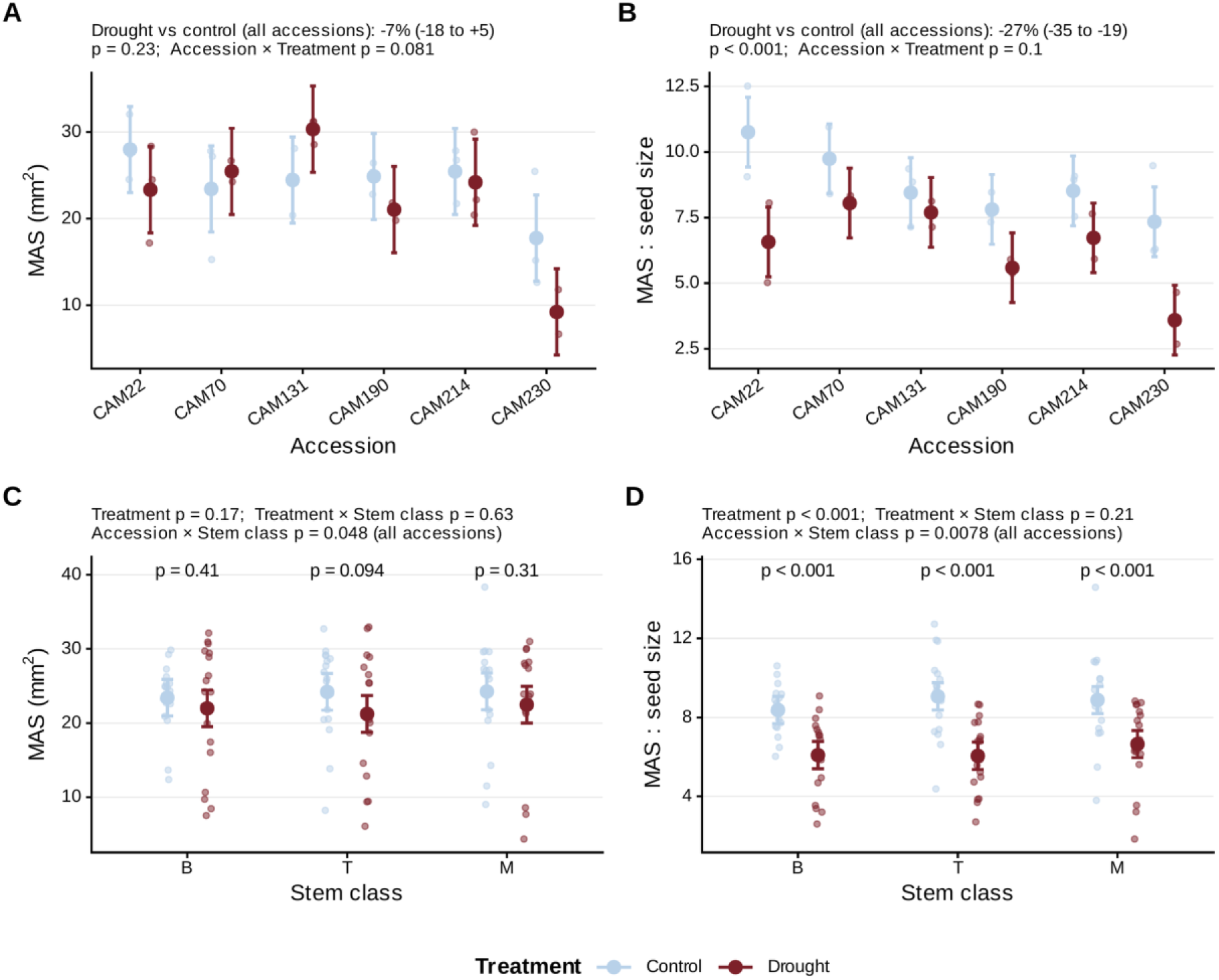
Seed mucilage of progeny from control and drought-grown plants, by accession and by stem class. **(A)** Mucilage area per seed (MAS) and **(B)** MAS : seed size ratio per accession. (C) MAS and **(D)** MAS : seed size ratio per stem class (B, basal secondary stem; T, top secondary stem; M, main stem main inflorescence), averaged over accessions. Large points and bars are estimated marginal means with 95% CI from linear mixed models (plant, and in C–D stem within plant, as random effects); small points are plant means (A–B) or stem means (C–D); n = 3 plants per accession and treatment. Subtitles give the drought effect averaged over accessions and the main interaction tests; P values in C–D are Holm-adjusted drought vs control contrasts.

### Drought shifts in non-adherent mucilage differ among accessions for Rha and Gal

The two mucilage fractions differed in composition (**Figure 4A**). Non-adherent mucilage contained more Rha than adherent mucilage (40.1 vs 26.3 molar %; P < 0.001) and slightly more Xyl (0.38 vs 0.25 molar %; P < 0.001), whereas adherent mucilage was richer in Gal (24.6 vs 13.3 molar %; P < 0.001), Ara (1.8 vs 0.3 molar %; P < 0.001), Glc (2.0 vs 0.2 molar %; P < 0.001) and Man (0.34 vs 0.24 molar %; P < 0.001). GalA was the most abundant monosaccharide in both fractions and did not differ between them (44.6 and 45.4 molar %; P = 0.42). Drought changed the composition of both mucilage fractions (**Figure 4B**). Monosaccharide composition also differed among accessions, for Rha, Ara, Gal and GalA in non-adherent mucilage and for Gal and Glc in adherent mucilage (P ≤ 0.038). Accession was the largest source of variation in the multivariate sugar profile of both fractions (PERMANOVA, R² = 0.44 non-adherent and 0.31 adherent, P ≤ 0.002). Maternal drought additionally shifted the composition of non-adherent mucilage (R² = 0.14, P < 0.001) and less strongly, of adherent mucilage (R² = 0.06, P = 0.046), and in both fractions this shift differed among accessions (A × T, P = 0.049 non-adherent and P = 0.041 adherent; **Figure S5**). Averaged over accessions, drought increased Rha (+2.1 molar %, +5%; P = 0.002, FDR q = 0.021) and Man (+0.12 molar %, +67%; P = 0.006, q = 0.026) in non-adherent mucilage (**Figure 4B**). The Rha increase depended on accession (A × T, P < 0.001) and was confined to CAM230 (+10.1 molar %, Holm P < 0.001; **Figure 4C**). Gal in non-adherent mucilage did not change on average (q = 0.13), but its response also depended on accession (A × T, P = 0.004): it decreased only in CAM230, from 10.5 to 1.6 molar % (Holm P = 0.002; **Figure 4D**). Ara in non-adherent mucilage, by contrast, did not change in CAM230 (0.24 vs 0.23 molar %; Holm P = 1.00; **Figure S6**), so drought lowered Gal without lowering Ara in this accession. In adherent mucilage, drought increased Glc (from 0.9 to 3.1 molar %; P = 0.003, q = 0.021) and decreased GalA (−5.5 molar %, −12%; P = 0.013, q = 0.044). The Glc increase also depended on accession (A × T, P = 0.007) and was significant only in CAM230 (+7.9 molar %, Holm P < 0.001; **Figure 4E**). The drought response differed between the two fractions for Gal, Glc and Man (treatment × fraction, P ≤ 0.010).

**Figure 4.**
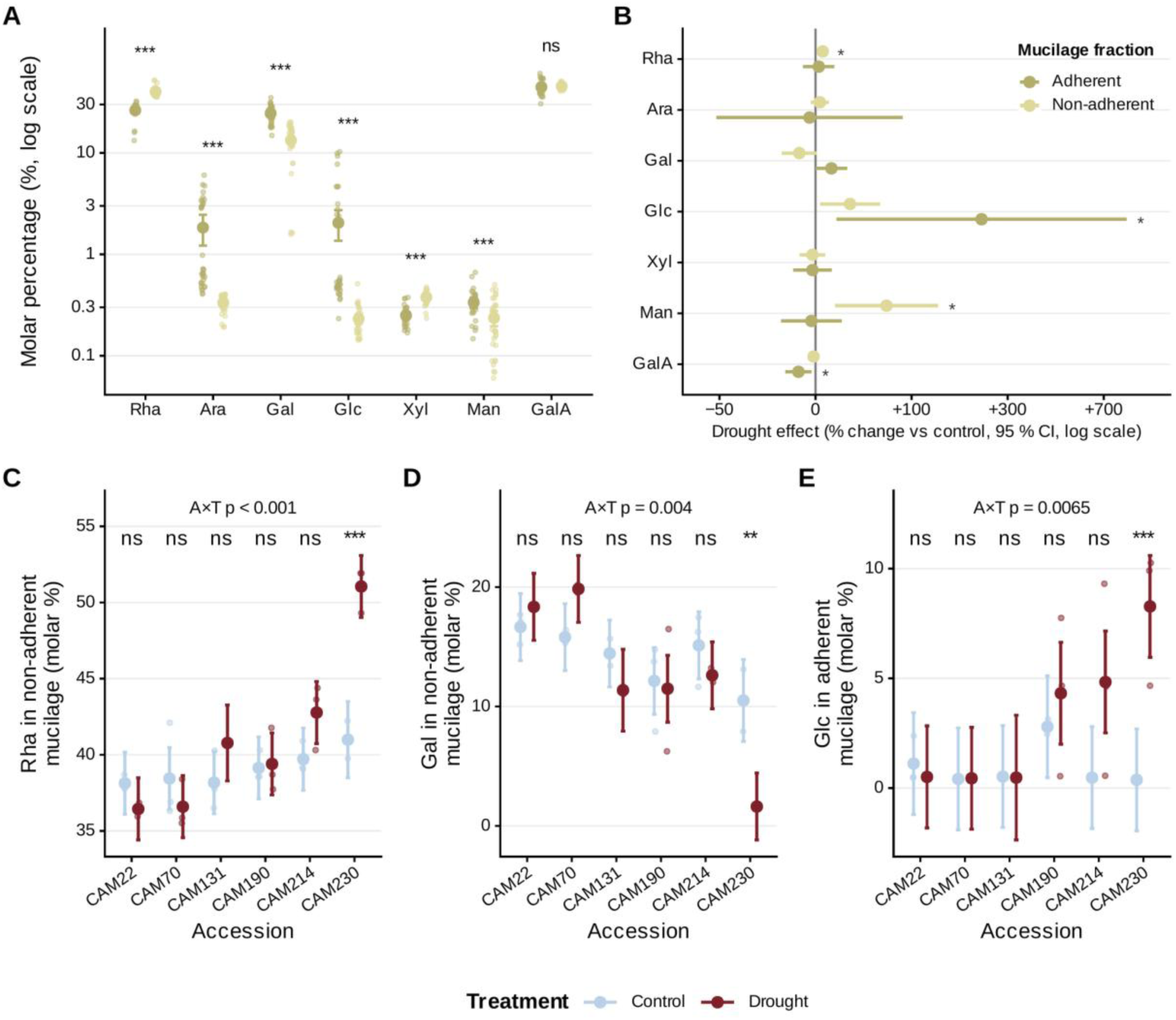
Monosaccharide composition of adherent and non-adherent seed mucilage from control and drought-grown plants. **(A)** Molar percentage of seven monosaccharides in adherent and non-adherent mucilage, averaged over accessions and treatments (log scale). Stars indicate differences between fractions, tested on the paired non-adherent minus adherent difference per plant (FDR-adjusted across monosaccharides). **(B)** Drought effect on each monosaccharide in each fraction, averaged over accessions (% change from control with 95% CI, log scale; stars, FDR-adjusted). **(C)** Rha and **(D)** Gal in non-adherent mucilage and **(E)** Glc in adherent mucilage per accession, the three cases with a significant accession × treatment interaction (A × T); stars are Holm-adjusted drought vs control contrasts within each accession. Large points and bars are estimated marginal means with 95% CI from linear models fitted per monosaccharide and fraction (A × T, one value per plant); small points are plant means of up to three capsule samples. Rha, rhamnose; Ara, arabinose; Gal, galactose; Glc, glucose; Xyl, xylose; Man, mannose; GalA, galacturonic acid. * P < 0.05, ** P < 0.01, *** P < 0.001; ns, not significant.

### Maternal drought leaves an intergenerational imprint on progeny germination under salt and osmotic stress

We next assessed whether maternal drought affected progeny germination under salt (NaCl) and osmotic (PEG) stress. Germination of 30 seeds per plant was scored daily for 3 days in water, in NaCl at −0.6 and −1.2 MPa, and in PEG at −0.4 to −1.0 MPa (**Figure 5A**). In water and PEG −0.4 MPa, almost all seeds germinated in both treatments (≥ 97% by day 3), whereas no seed germinated in PEG −0.8 or −1.0 MPa. In water, maternal drought slowed germination (day 1: 92% vs 97%; OR = 0.24, 95% CI 0.08–0.71, P = 0.009), but final germination was complete in both treatments (99.8% and 100%). Maternal drought reduced final germination under moderate salt and osmotic stress (P < 0.001; **Figure 5B**). Progeny of drought-stressed plants had lower odds of germinating under NaCl −0.6 MPa (92% vs 98%; OR = 0.30, 95% CI 0.14-0.64, P = 0.002), NaCl −1.2 MPa (87% vs 93%; OR = 0.46, 0.27-0.81, P = 0.007) and PEG −0.6 MPa (21% vs 35%; OR = 0.42, 0.27-0.65, P < 0.001). The reduction was similar across the three stress conditions (treatment × condition, P = 0.62). At the same water potential (−0.6 MPa), PEG was far more inhibitory than NaCl, with odds of germinating 70 to 99 times higher in NaCl than in PEG (P < 0.001). Furthermore, the A × T interaction was significant only under PEG −0.6 MPa (P = 0.006; NaCl −0.6 MPa P = 0.67; NaCl −1.2 MPa P = 0.079; **Figure 5C**), where germination of progeny from drought-grown plants fell from 29% to 7% in CAM70 (OR = 0.17), from 31% to 8% in CAM131 (OR = 0.18) and from 39% to 13% in CAM230 (OR = 0.24; Holm-adjusted P = 0.003 for each), whereas CAM22, CAM190 and CAM214 were not affected (Holm-adjusted P ≥ 0.42). In CAM70, CAM131 and CAM230, germination of drought progeny did not catch up with that of control progeny by day 3, indicating reduced rather than merely delayed germination within the assay period (**Figure 5A, C**). A delay was instead suggested for CAM70 under NaCl −0.6 MPa: on day 1, 3% of seeds from drought-stressed plants had germinated compared with 38% of seeds from control plants, but by day 3 germination was similar (92% vs 99%). This contrast was not significant after Holm adjustment (OR = 0.04, P = 0.11). Finally, we tested whether seed traits and mucilage were associated with progeny germination (**Table S3**). Seed size and seed weight were not associated with germination (FDR-adjusted P ≥ 0.18) nor with mucilage composition (FDR-adjusted P ≥ 0.08). Within accessions and treatments, two mucilage traits were associated with germination, both only under NaCl −1.2 MPa. Among accessions, the drought change in germination tracked neither the drought change in MAS (Spearman ρ = −0.44 to 0.26, P ≥ 0.38) nor that in any monosaccharide (FDR-adjusted P ≥ 0.83). Specific mucilage traits were therefore associated with germination within accessions, but did not explain the drought-induced loss of germination among accessions.

**Figure 5.**
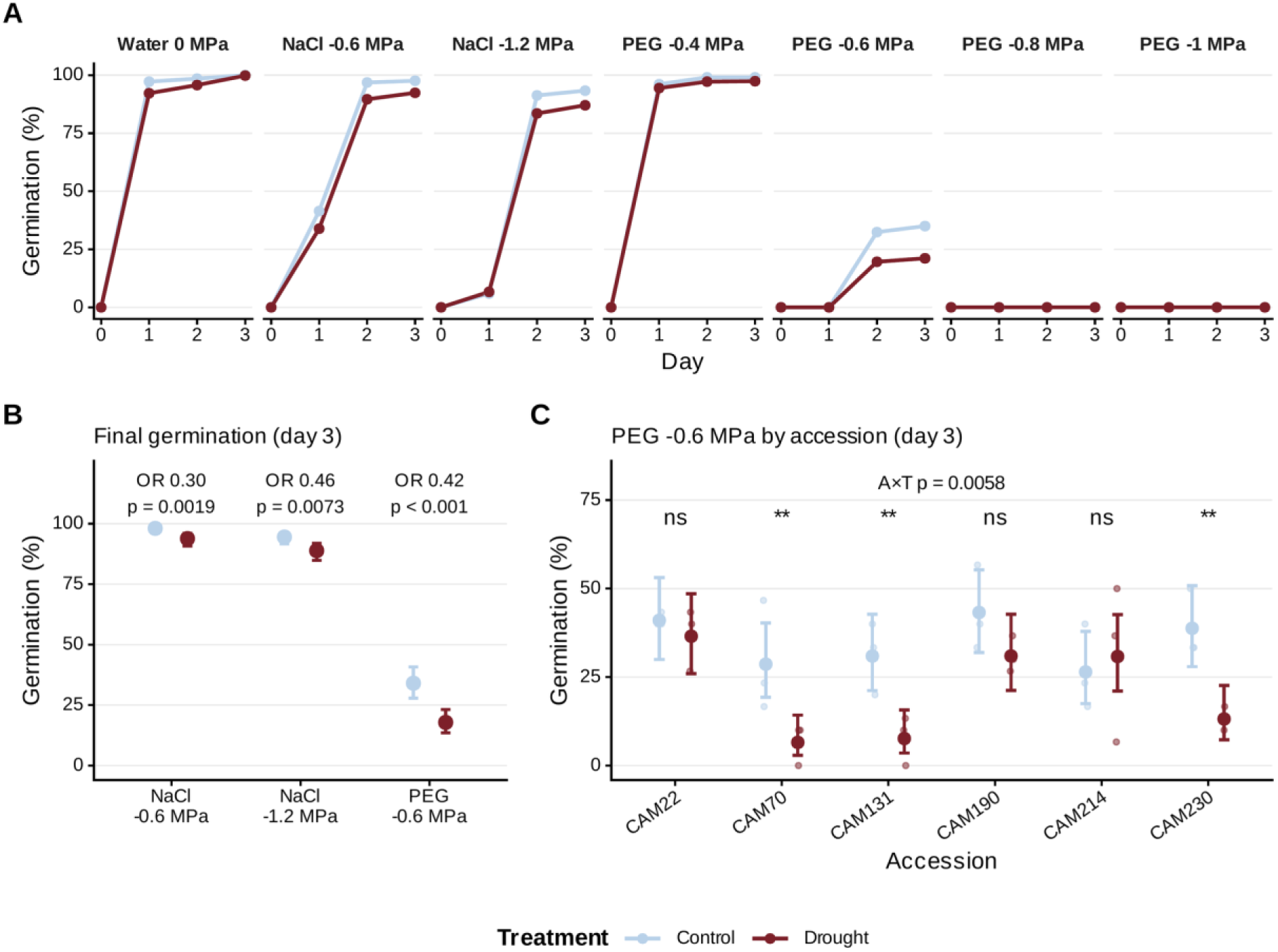
Germination of progeny seeds from control and drought-grown *Camelina* plants under osmotic (PEG) and salt (NaCl) stress. **(A)** Observed germination over days 0-3 in water and at each water potential, pooled over six accessions (18 plants per treatment, 30 seeds per plant**). (B)** Final germination (day 3) under NaCl −0.6, NaCl −1.2 and PEG −0.6 MPa, averaged over accessions (estimated marginal means from a binomial GLMM); OR, odds ratio of germination, drought vs control. **(C)** Final germination under PEG −0.6 MPa per accession (estimated marginal means from a binomial GLMM); small dots are individual plants; stars give the Holm-adjusted drought vs control contrast within each accession (** P < 0.01; ns, not significant); A × T, likelihood-ratio test of the accession × treatment interaction. Error bars are 95 % CI.

## Discussion

The main objective of this study was to determine how drought affects *Camelina* and whether these effects carry over to the next generation as adaptive traits. Drought affected every level we measured, from leaf physiology to the germination of the progeny, and most responses were consistent among the six accessions (**Figure 6**). Accessions differed in the dynamics of shoot growth, in the drought response of mucilage composition and in germination under PEG. At the same time, mucilage area scaled with seed size within each treatment but did not increase when drought made seeds larger. Within accessions and treatments, two mucilage traits (mucilage area and glucose in adherent mucilage) were associated with germination, but only under strong salt stress (NaCl −1.2 MPa). Among accessions, neither the seed nor the mucilage traits explained the lower germination of drought progeny, which does not support an adaptive role of these traits under abiotic stress.

**Figure 6.**
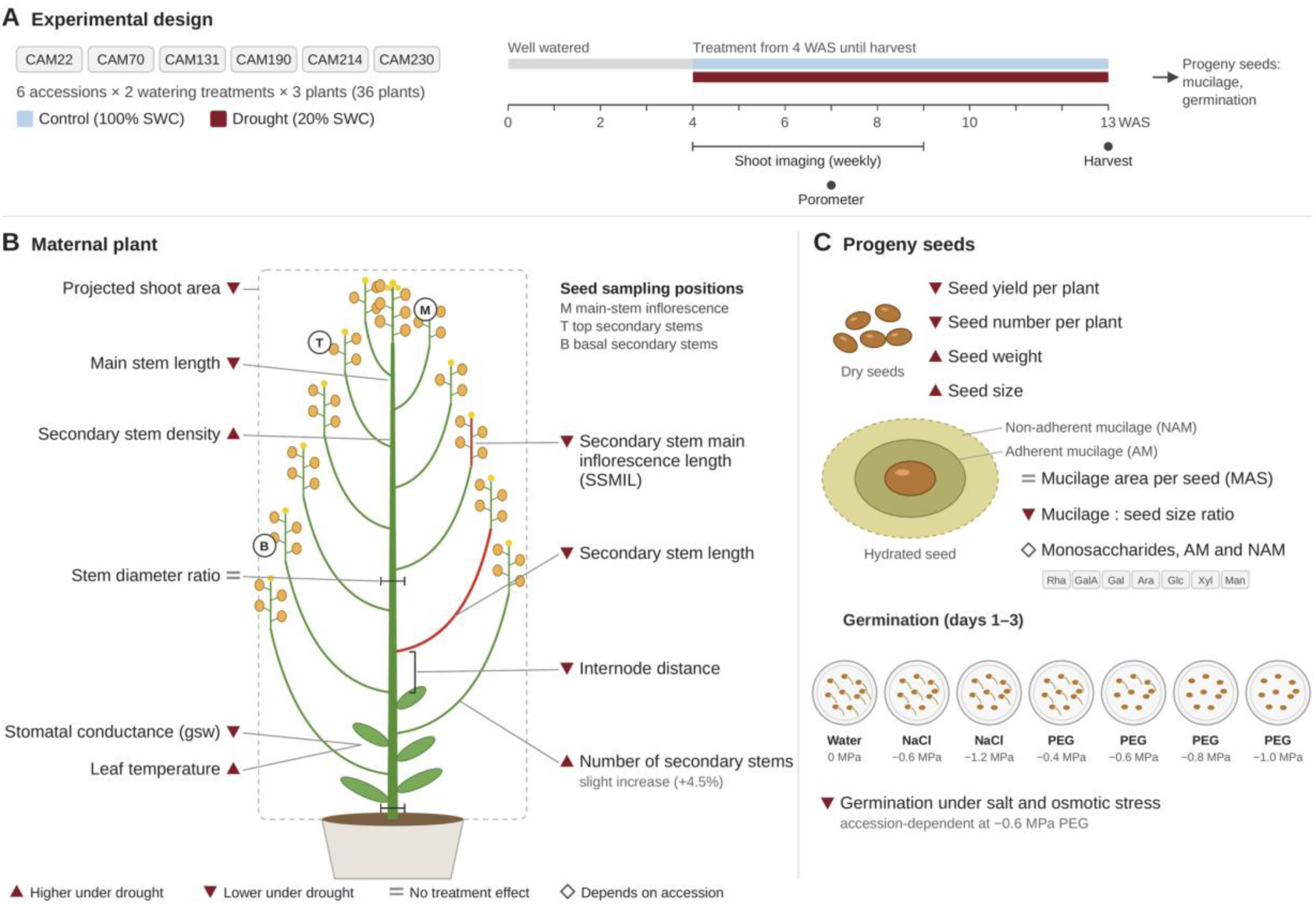
Overview of the drought responses of maternal *Camelina* plants and their progeny seeds. **(A)** Experimental design. **(B)** Maternal plant and **(C)** progeny seed traits; symbols give the drought effect averaged over accessions: ▴ higher and ▾ lower under drought, = no effect of treatment, ◇ drought effect differs among accessions. M, T and B mark the seed sampling positions (main stem main inflorescence, top and basal secondary stems), and red lines mark the measured secondary stem and SSMIL. AM, adherent mucilage; MAS, mucilage area per seed; NAM, non-adherent mucilage; SSMIL, secondary stem main inflorescence length; SWC, soil water content; WAS, weeks after sowing.

### Drought-stressed mother plants cope by closing their stomata and building a compact shoot

Drought-stressed plants closed their stomata in all accessions. Stomatal closure is the typical first response of plants to soil drying: it limits water loss but also restricts CO₂ uptake and photosynthesis, and species differ in how tightly they regulate it (Chaves et al., 2003; Tardieu and Simonneau, 1998). In four *Camelina* lines exposed to drought from the onset of flowering, stomatal conductance and net photosynthesis fell by about 80% and 60%, and the lines differed in the size of this response (Alberghini et al., 2025). Here, stomatal conductance fell by 95% and leaf temperature rose by 1.3 °C in all six accessions, without detectable differences among them. Accordingly, shoot growth was reduced in all accessions, but the reduction persisted only in some. Reduced growth is a common consequence of drought, reported also in *Camelina* (Waraich et al., 2017) and other Brassicaceae (Hasanuzzaman, 2020), and the observed treatment convergence could at least partly reflect the decline of control plants rather than a recovery of drought-stressed plants. Drought-stressed plants also built a more compact shoot in all accessions, reducing the elongation of stems rather than their number. Reduced inflorescence stem growth under water limitation has also been described in *Arabidopsis* (Su et al., 2013), and in sorghum drought reduced internode elongation without changing internode diameter (Perrier et al., 2017). In *Camelina* lines exposed to drought from the onset of flowering, plants were 15% shorter, lateral branch biomass was halved and silicle number fell by 26% (Alberghini et al., 2025), and in genome-edited flowering-time mutants of *Camelina*, drought altered branching and yield, whereas flowering time changed little (Vayssières et al., 2026).

### Drought-stressed mother plants prioritize individual seeds, but the effect on the progeny is not adaptive

Drought-stressed plants produced fewer but larger and heavier seeds, consistently across accessions. Theory predicts that a mother plant should keep the size of its offspring near an optimum and adjust their number to the resources available (Smith and Fretwell, 1974), and in most grain crops, such as wheat and soybean, seed number is indeed more plastic than seed size (Sadras, 2007). Under water deficit, seed number is reduced through the abortion of the youngest reproductive organs, which allows older organs to complete their development when resources are limited (Turc and Tardieu, 2018), and drought may have reduced seed number in the same way here. Seed size, however, did not remain constant but increased by 28%, and seed weight by 38%. In *Camelina* lines exposed to drought only from the onset of flowering, seed weight also increased, but by just 5%, and seed number was not reduced on average (Alberghini et al., 2025); across 54 spring accessions grown at three field sites, seed weight was stable (Silvestre et al., 2026). The stronger response found here may reflect the earlier onset of drought, before bolting, which would allow the plant to adjust seed number before seed filling, but this remains to be tested by varying the timing of drought. Larger seeds often improve seedling performance under stressful conditions (Leishman et al., 2000; Lloret et al., 1999). In a previous study in *Camelina*, the effect of seed size on germination under PEG depended on biotype: large-seeded spring types germinated best, whereas among winter types small-seeded ones performed best (Čanak et al., 2020). In our study, PEG inhibited germination far more than NaCl at the same water potential and the larger seeds of drought-stressed plants germinated less. Germination of *Camelina* is relatively robust to salt: among 57 accessions, most still germinated to at least 90% at 200 mM NaCl, although germination was delayed and accessions differed in how fast and how uniformly they germinated (Mastroberardino et al., 2025).

Seed mucilage is considered a substantial carbon investment of the mother plant. *Arabidopsis* mutants that lack mucilage (*gl2*, *mum4*) accumulate more seed oil, which led to the hypothesis that mucilage and oil compete for photosynthate, although loss of mucilage alone did not raise oil content in every mutant (Shi et al., 2012). If such competition had shaped the drought response, drought-stressed plants would have produced larger seeds at the expense of mucilage. However, drought-stressed plants maintained the mucilage area per seed while increasing seed size, so mucilage area only decreased relative to seed size. Our results therefore do not support a trade-off between seed size and mucilage production. Mucilage is produced by the epidermal cells of the seed coat, a tissue of maternal origin (Arsovski et al., 2010; Haughn and Western, 2012), and its extent appears to respond less to the maternal water supply than seed filling does. Mucilage did not explain the lower germination either: at equal seed size, mucilage area decreased as much in CAM22 (60%), whose germination under PEG was not reduced, as in CAM230 (58%), whose germination was. Seed mucilage can improve water uptake and germination under salt and osmotic stress in some species (Bhatt et al., 2016; Chen et al., 2018; Mira et al., 2023; Sohrabizadeh et al., 2023), and in four *Plantago* species, seeds with more mucilage germinated at lower water potentials and produced seedlings that survived water deficit better (Teixeira et al., 2020). In *Camelina*, however, the lower relative mucilage of drought progeny does not appear to be the main cause of their lower germination.

Maternal drought shifted mucilage composition mainly in the non-adherent fraction and most strongly in CAM230. The observed contrast between Ara and Gal was of particular interest, because in *Camelina* these sugars make up a larger part of mucilage than in *Arabidopsis*, and vary strongly among accessions in non-adherent mucilage. We have previously observed a strong genetic association between these sugars, which suggests variation in arabinogalactan-type polymers. Maternal drought uncoupled the two sugars in CAM230, Gal in non-adherent mucilage fell from 10.5 to 1.6 molar %, while Ara did not change. Gal-containing side chains matter for the properties of mucilage. In *Arabidopsis*, the β-galactosidase MUM2 removes galactose and galactan branches from rhamnogalacturonan I, which increases mucilage swelling and extrusion on hydration (Macquet et al., 2007b), and natural *mum2* variants produce seeds that float, a trait proposed, though not demonstrated, to aid dispersal by water (Saez-Aguayo et al., 2014). Changes in mucilage polysaccharides also alter the adherence and organization of mucilage (Griffiths et al., 2014). It remains to be tested whether a drought-induced loss of Gal could change mucilage hydration, release or seed buoyancy in *Camelina*. Drought also raised Man, a component of mucilage heteromannan, in non-adherent mucilage. In *Arabidopsis*, *muci10* mutants, whose mucilage heteromannan is less galactosylated and which have less Gal and Man in mucilage, germinate better under salt, especially CaCl₂, but not under purely osmotic stress (Yang et al., 2021). Across our accessions, however, the composition shifts did not follow the germination losses under salt or PEG; CAM70 and CAM131, for example, lost germination under PEG without the shifts seen in CAM230. Fitness measures such as seedling establishment and survival in soil, seed dispersal and persistence, and seed production of the progeny, tested along a gradient of drought intensities and in the field, could contribute to better understand plasticity in *Camelina* (Murren et al., 2015). In soil, mucilage may also contribute to seed microbiome assembly and microbial interactions in the spermosphere (Nelson, 2004; Schiltz et al., 2015; Yang et al., 2012), so that drought-induced changes in mucilage, such as the loss of Gal in CAM230, could alter which microorganisms the progeny recruits.

### Accessions differ in trait values, but their drought responses differed only in growth dynamics, mucilage composition and germination under PEG

*Camelina* is considered to have low genetic diversity (Brock et al., 2024), yet it shows considerable plasticity in response to stress: across 54 spring accessions grown at three field sites, yield and oil content were sensitive to genotype × environment interactions, whereas thousand-seed weight was stable (Silvestre et al., 2026), and four lines differed in their physiological, yield and oil responses to drought (Alberghini et al., 2025). In our study, the accessions that differed in their drought response were not the same for growth of the mother plant, mucilage composition and progeny germination, which suggests that they do not share a single cause. Because plasticity and trait means can have partly separate genetic bases and the evolution of plasticity depends on the genetic variation available (Laitinen and Nikoloski, 2019; Murren et al., 2015), these accession-specific responses are the most promising starting points for genetic mapping and breeding in larger panels, as has been done for germination under salt in 211 spring accessions (Luo et al., 2020). Moreover, accession explained more of the variation in mucilage composition than drought did (44% vs 14% in non-adherent and 31% vs 6% in adherent mucilage). The two mucilage fractions also differed consistently, with more Rha and Xyl in non-adherent mucilage and more Gal, Ara, Glc and Man in adherent mucilage, which resembles *Arabidopsis*. In *Arabidopsis*, both layers consist largely of rhamnogalacturonan I, but differ in composition and structure (Macquet et al., 2007a; Voiniciuc et al., 2016), with an extensive natural variation among accessions (Voiniciuc et al., 2016), whereas the composition of the outer, non-adherent layer varies little among more than 300 accessions (Fabrissin et al., 2019), unlike the strong accession effect on non-adherent mucilage found here. Our conclusions rest on three plants per accession and treatment, which limits the power to detect differences among accessions; a non-significant A × T interaction indicates no evidence that accessions differ, not evidence that they respond alike.

## Conclusions

Drought affected *Camelina* from leaf physiology to the next generation. Drought-stressed plants developed a more compact shoot and prioritized the provisioning of individual seeds, producing fewer but larger, heavier seeds. This priority did not extend to the seed coat: the larger seeds had a smaller mucilage area relative to their size and germinated less under salt and osmotic stress, so larger seeds were not better prepared for stressful germination conditions, indicating a cost of maternal drought for the progeny rather than an adaptive response. These responses were mostly consistent among the six accessions, whereas accessions differed in the dynamics of shoot growth, in the monosaccharide composition of mucilage and in germination under PEG. Breeding for drought-prone, low-input systems should consider effects on the progeny alongside yield.

## Conflict of interest

The authors declare that the research was conducted in the absence of any commercial or financial relationships that could be construed as a potential conflict of interest.

## Author contributions

**MBA:** Conceptualization, Data curation, Formal analysis, Investigation, Methodology, Software, Visualization, Writing – original draft, Writing – review & editing. **AGO:** Investigation, Methodology, Writing – review & editing. **JK:** Investigation, Writing – review & editing. **SS:** Investigation, Writing – review & editing. **BU:** Conceptualization, Funding acquisition, Methodology, Project administration, Resources, Supervision, Writing – review & editing. **CL:** Conceptualization, Formal analysis, Methodology, Supervision, Visualization, Writing – original draft, Writing – review & editing. All authors read and approved the submitted version.

## Funding

This work was funded by the Deutsche Forschungsgemeinschaft (DFG, German Research Foundation) through the Collaborative Research Centre TRR 341 Plant Ecological Genetics (project number 456082119) and by the European Union’s Horizon 2020 research and innovation programme through the UNTWIST project (grant agreement No. 862524). **BU** is also funded by the DFG under Germany’s Excellence Strategy, EXC 2048/1, Project ID 390686111.

## Acknowledgments

We thank the gardeners of Heinrich-Heine-Universität Düsseldorf for providing and maintaining the greenhouse facilities. We also thank Barbara Schulten for her help with maintaining the plants in the greenhouse, taking plant measurements and preparing the ion chromatography samples. During the preparation of this manuscript, the authors used Claude (model Claude Opus 5.5; Anthropic, https://claude.ai) to review the draft, check the statistical analyses and cited references, and draw the schematic figures. The authors reviewed and edited all output and take full responsibility for the content of this publication.

## Supplementary material

The Supplementary Material (Tables S1–S3 and Figures S1–S6) is provided as a separate file.

## Data availability statement

The raw data, analysis scripts and model outputs supporting the conclusions of this study are available as an Annotated Research Context (ARC) on the DataPLANT DataHUB: https://doi.org/10.60534/weahs-c3h09.

## Notes

https://doi.org/10.60534/weahs-c3h09

### Competing Interest Statement

The authors have declared no competing interest.

## References

Aitchison, J. (1986). The Statistical Analysis of Compositional Data. London: Chapman and Hall.

Alberghini, B., Vicino, M., Zanetti, F., Silvestre, S., Haslam, R., Zegada-Lizarazu, W., et al. (2025). Assessing different physiological, seed yield and quality responses of camelina lines to drought. Ind. Crops Prod. 234:121528. doi: 10.1016/j.indcrop.2025.121528

Anderson, M. J. (2001). A new method for non-parametric multivariate analysis of variance. Austral Ecol. 26, 32–46. doi: 10.1111/j.1442-9993.2001.01070.pp.x

Arsovski, A. A., Haughn, G. W., and Western, T. L. (2010). Seed coat mucilage cells of *Arabidopsis thaliana* as a model for plant cell wall research. Plant Signal. Behav. 5, 796–801. doi: 10.4161/psb.5.7.11773

Basu, S., Ramegowda, V., Kumar, A., and Pereira, A. (2016). Plant adaptation to drought stress. F1000Res. 5:1554. doi: 10.12688/f1000research.7678.1

Bates, D., Mächler, M., Bolker, B. M., and Walker, S. C. (2015). Fitting linear mixed-effects models using lme4. J. Stat. Softw. 67, 1–48. doi: 10.18637/jss.v067.i01

Benavides-Acevedo, M. (2025a). Fiji macro to count and measure the area of *Camelina sativa* seeds. Zenodo. doi: 10.5281/zenodo.17937951

Benavides-Acevedo, M. (2025b). Fiji/ImageJ macro for quantitative analysis of *Camelina sativa* seed mucilage from segmented images. Zenodo. doi: 10.5281/zenodo.17939990

Benjamini, Y., and Hochberg, Y. (1995). Controlling the false discovery rate: a practical and powerful approach to multiple testing. J. R. Stat. Soc. Series B Stat. Methodol. 57, 289–300. doi: 10.1111/j.2517-6161.1995.tb02031.x

Bhatt, A., Santo, A., and Gallacher, D. (2016). Seed mucilage effect on water uptake and germination in five species from the hyper-arid Arabian desert. J. Arid Environ. 128, 73–79. doi: 10.1016/j.jaridenv.2016.01.008

Brock, J. R., Bird, K. A., Platts, A. E., Gomez-Cano, F., Gupta, S. K., Palos, K., et al. (2024). Exploring genetic diversity, population structure, and subgenome differences in the allopolyploid *Camelina sativa*: implications for future breeding and research studies. Hortic. Res. 11:uhae247. doi: 10.1093/hr/uhae247

Brooks, M. E., Kristensen, K., van Benthem, K. J., Magnusson, A., Berg, C. W., Nielsen, A., et al. (2017). glmmTMB balances speed and flexibility among packages for zero-inflated generalized linear mixed modeling. R J. 9, 378–400. doi: 10.32614/RJ-2017-066

Čanak, P., Jeromela, A. M., Vujošević, B., Kiprovski, B., Mitrović, B., Alberghini, B., et al. (2020). Is drought stress tolerance affected by biotypes and seed size in the emerging oilseed crop Camelina? Agronomy 10:1856. doi: 10.3390/agronomy10121856

Chaves, M. M., Maroco, J. P., and Pereira, J. S. (2003). Understanding plant responses to drought: from genes to the whole plant. Funct. Plant Biol. 30, 239–264. doi: 10.1071/FP02076

Chen, Y., Zhang, L., Lu, X., Lan, X., Shen, M., and Lu, C. (2018). Role of mucilage during achene germination and sprout growth of the endangered Tibetan medicinal herb *Mirabilis himalaica* (Nyctaginaceae) exposed to abiotic stresses. J. Plant Ecol. 11, 328–337. doi: 10.1093/jpe/rtx047

Chung, Y., Rabe-Hesketh, S., Dorie, V., Gelman, A., and Liu, J. (2013). A nondegenerate penalized likelihood estimator for variance parameters in multilevel models. Psychometrika 78, 685–709. doi: 10.1007/s11336-013-9328-2

Donohue, K. (2009). Completing the cycle: maternal effects as the missing link in plant life histories. Philos. Trans. R. Soc. Lond. B Biol. Sci. 364, 1059–1074. doi: 10.1098/rstb.2008.0291

Engelbrecht, M., Bochet, E., and García-Fayos, P. (2014). Mucilage secretion: an adaptive mechanism to reduce seed removal by soil erosion? Biol. J. Linn. Soc. 111, 241–251. doi: 10.1111/bij.12198

Fabrissin, I., Cueff, G., Berger, A., Granier, F., Sallé, C., Poulain, D., et al. (2019). Natural variation reveals a key role for rhamnogalacturonan I in seed outer mucilage and underlying genes. Plant Physiol. 181, 1498–1518. doi: 10.1104/pp.19.00763

Food and Agriculture Organization of the United Nations (2023). The impact of disasters on agriculture and food security. https://www.fao.org/newsroom/detail/first-ever-global-estimation-of-the-impact-of-disasters-on-agriculture/en/

Fox, J., and Weisberg, S. (2019). An R Companion to Applied Regression, 3rd Edn. Thousand Oaks, CA: Sage.

Griffiths, J. S., Tsai, A. Y. L., Xue, H., Voiniciuc, C., Šola, K., Seifert, G. J., et al. (2014). SALT-OVERLY SENSITIVE5 mediates *Arabidopsis* seed coat mucilage adherence and organization through pectins. Plant Physiol. 165, 991–1004. doi: 10.1104/pp.114.239400

Guo, J., Du, M., Tian, H., and Wang, B. (2020). Exposure to high salinity during seed development markedly enhances seedling emergence and fitness of the progeny of the extreme halophyte *Suaeda salsa*. Front. Plant Sci. 11:1291. doi: 10.3389/fpls.2020.01291

Hartig, F. (2026). DHARMa: residual diagnostics for hierarchical (multi-level/mixed) regression models. R package version 0.5.0. doi: 10.32614/CRAN.package.DHARMa

Hasanuzzaman, M. (ed.) (2020). The Plant Family Brassicaceae: Biology and Physiological Responses to Environmental Stresses. Singapore: Springer. doi: 10.1007/978-981-15-6345-4

Hatzig, S. V., Nuppenau, J. N., Snowdon, R. J., and Schießl, S. V. (2018). Drought stress has transgenerational effects on seeds and seedlings in winter oilseed rape (*Brassica napus* L.). BMC Plant Biol. 18:297. doi: 10.1186/s12870-018-1531-y

Haughn, G. W., and Western, T. L. (2012). *Arabidopsis* seed coat mucilage is a specialized cell wall that can be used as a model for genetic analysis of plant cell wall structure and function. Front. Plant Sci. 3:64. doi: 10.3389/fpls.2012.00064

Herman, J. J., and Sultan, S. E. (2011). Adaptive transgenerational plasticity in plants: case studies, mechanisms, and implications for natural populations. Front. Plant Sci. 2:102. doi: 10.3389/fpls.2011.00102

Holm, S. (1979). A simple sequentially rejective multiple test procedure. Scand. J. Stat. 6, 65–70.

IPCC (2021). “Summary for policymakers,” in Climate Change 2021: The Physical Science Basis. Contribution of Working Group I to the Sixth Assessment Report of the Intergovernmental Panel on Climate Change, eds. V. Masson-Delmotte, P. Zhai, A. Pirani, S. L. Connors, C. Péan, S. Berger, et al. (Cambridge: Cambridge University Press), 3–32. doi: 10.1017/9781009157896.001

Kuznetsova, A., Brockhoff, P. B., and Christensen, R. H. B. (2017). lmerTest package: tests in linear mixed effects models. J. Stat. Softw. 82, 1–26. doi: 10.18637/jss.v082.i13

Laitinen, R. A. E., and Nikoloski, Z. (2019). Genetic basis of plasticity in plants. J. Exp. Bot. 70, 739–745. doi: 10.1093/jxb/ery404

Leishman, M. R., Wright, I. J., Moles, A. T., and Westoby, M. (2000). “The evolutionary ecology of seed size,” in Seeds: The Ecology of Regeneration in Plant Communities, 2nd Edn, ed. M. Fenner (Wallingford: CABI Publishing), 31–57.

Lenth, R. V., and Piaskowski, J. (2026). emmeans: estimated marginal means, aka least-squares means. R package version 2.0.4. doi: 10.32614/CRAN.package.emmeans

Lloret, F., Casanovas, C., and Peñuelas, J. (1999). Seedling survival of Mediterranean shrubland species in relation to root:shoot ratio, seed size and water and nitrogen use. Funct. Ecol. 13, 210–216. doi: 10.1046/j.1365-2435.1999.00309.x

Luo, Z., Szczepanek, A., and Abdel-Haleem, H. (2020). Genome-wide association study (GWAS) analysis of camelina seedling germination under salt stress condition. Agronomy 10:1444. doi: 10.3390/agronomy10091444

Macquet, A., Ralet, M. C., Kronenberger, J., Marion-Poll, A., and North, H. M. (2007a). In situ, chemical and macromolecular study of the composition of *Arabidopsis thaliana* seed coat mucilage. Plant Cell Physiol. 48, 984–999. doi: 10.1093/pcp/pcm068

Macquet, A., Ralet, M. C., Loudet, O., Kronenberger, J., Mouille, G., Marion-Poll, A., et al. (2007b). A naturally occurring mutation in an *Arabidopsis* accession affects a β-D-galactosidase that increases the hydrophilic potential of rhamnogalacturonan I in seed mucilage. Plant Cell 19, 3990–4006. doi: 10.1105/tpc.107.050179

Mastroberardino, R., Zanetti, F., and Monti, A. (2025). Exploring intraspecific variation in salinity tolerance at germination and seedling development stages in *Camelina sativa*. Front. Plant Sci. 16:1713651. doi: 10.3389/fpls.2025.1713651

Miart, F., Fontaine, J. X., Pineau, C., Demailly, H., Thomasset, B., Van Wuytswinkel, O., et al. (2018). MuSeeQ, a novel supervised image analysis tool for the simultaneous phenotyping of the soluble mucilage and seed morphometric parameters. Plant Methods 14:112. doi: 10.1186/s13007-018-0377-5

Miart, F., Fournet, F., Dubrulle, N., Petit, E., Demailly, H., Dupont, L., et al. (2019). Cytological approaches combined with chemical analysis reveals the layered nature of flax mucilage. Front. Plant Sci. 10:684. doi: 10.3389/fpls.2019.00684

Mira, S., Veiga-Barbosa, L., Martínez-Laborde, J. B., Pérez-García, F., and González-Benito, M. E. (2023). Effect of osmotic stress and salinity on germination and mucilage expansion of seeds of *Plantago albicans* L. (Plantaginaceae): inter-population variation. Plant Species Biol. 38, 286–297. doi: 10.1111/1442-1984.12430

Mueller-Linow, M. (2022). A versatile tool for semantic plant segmentation via color space thresholding. Zenodo. doi: 10.5281/zenodo.6443757

Murren, C. J., Auld, J. R., Callahan, H., Ghalambor, C. K., Handelsman, C. A., Heskel, M. A., et al. (2015). Constraints on the evolution of phenotypic plasticity: limits and costs of phenotype and plasticity. Heredity 115, 293–301. doi: 10.1038/hdy.2015.8

Nelson, E. B. (2004). Microbial dynamics and interactions in the spermosphere. Annu. Rev. Phytopathol. 42, 271–309. doi: 10.1146/annurev.phyto.42.121603.131041

Nicotra, A. B., Atkin, O. K., Bonser, S. P., Davidson, A. M., Finnegan, E. J., Mathesius, U., et al. (2010). Plant phenotypic plasticity in a changing climate. Trends Plant Sci. 15, 684–692. doi: 10.1016/j.tplants.2010.09.008

North, H. M., Berger, A., Saez-Aguayo, S., and Ralet, M.-C. (2014). Understanding polysaccharide production and properties using seed coat mutants: future perspectives for the exploitation of natural variants. Ann. Bot. 114, 1251–1263. doi: 10.1093/aob/mcu011

Oksanen, J., Simpson, G. L., Blanchet, F. G., Kindt, R., Legendre, P., Minchin, P. R., et al. (2026). vegan: community ecology package. R package version 2.7–5. doi: 10.32614/CRAN.package.vegan

Pedersen, T. L. (2025). patchwork: the composer of plots. R package version 1.3.2. https://patchwork.data-imaginist.com

Penfield, S., and MacGregor, D. R. (2017). Effects of environmental variation during seed production on seed dormancy and germination. J. Exp. Bot. 68, 819–825. doi: 10.1093/jxb/erw436

Perera, S. P., McIntosh, T., Coutu, C., Tyler, R. T., Hegedus, D. D., and Wanasundara, J. P. D. (2022). Profiling and characterization of *Camelina sativa* (L.) Crantz meal proteins. J. Am. Oil Chem. Soc. 99, 873–889. doi: 10.1002/aocs.12634

Perrier, L., Rouan, L., Jaffuel, S., Clément-Vidal, A., Roques, S., Soutiras, A., et al. (2017). Plasticity of sorghum stem biomass accumulation in response to water deficit: a multiscale analysis from internode tissue to plant level. Front. Plant Sci. 8:1516. doi: 10.3389/fpls.2017.01516

Pinheiro, J. C., and Bates, D. M. (2000). Mixed-Effects Models in S and S-PLUS. New York: Springer. doi: 10.1007/b98882

R Core Team (2026). R: a language and environment for statistical computing. Version 4.5.3. Vienna: R Foundation for Statistical Computing. https://www.R-project.org/

Ralet, M.-C., Crépeau, M.-J., Vigouroux, J., Tran, J., Berger, A., Sallé, C., et al. (2016). Xylans provide the structural driving force for mucilage adhesion to the *Arabidopsis* seed coat. Plant Physiol. 171, 165–178. doi: 10.1104/pp.16.00211

Sadras, V. O. (2007). Evolutionary aspects of the trade-off between seed size and number in crops. Field Crops Res. 100, 125–138. doi: 10.1016/j.fcr.2006.07.004

Saez-Aguayo, S., Rondeau-Mouro, C., Macquet, A., Kronholm, I., Ralet, M. C., Berger, A., et al. (2014). Local evolution of seed flotation in *Arabidopsis*. PLoS Genet. 10:e1004221. doi: 10.1371/journal.pgen.1004221

Schiltz, S., Gaillard, I., Pawlicki-Jullian, N., Thiombiano, B., Mesnard, F., and Gontier, E. (2015). A review: what is the spermosphere and how can it be studied? J. Appl. Microbiol. 119, 1467–1481. doi: 10.1111/jam.12946

Schindelin, J., Arganda-Carreras, I., Frise, E., Kaynig, V., Longair, M., Pietzsch, T., et al. (2012). Fiji: an open-source platform for biological-image analysis. Nat. Methods 9, 676–682. doi: 10.1038/nmeth.2019

Shi, L., Katavic, V., Yu, Y., Kunst, L., and Haughn, G. (2012). *Arabidopsis glabra2* mutant seeds deficient in mucilage biosynthesis produce more oil. Plant J. 69, 37–46. doi: 10.1111/j.1365-313X.2011.04768.x

Silvestre, S., Prigent, S., Pétriacq, P., Hassall, K. L., Le Boulch, M., Da Costa, A., et al. (2026). Assessment of phenotypic trait plasticity in the oilseed *Camelina sativa* using integrated early stage abiotic stress and field studies. Plant Physiol. 200:kiag052. doi: 10.1093/plphys/kiag052

Smith, C. C., and Fretwell, S. D. (1974). The optimal balance between size and number of offspring. Am. Nat. 108, 499–506. doi: 10.1086/282929

Sohrabizadeh, Z., Sodaeizadeh, H., and Mosleh Arani, A. (2023). Role of mucilage in improving seedling emergence of *Ocimum basilicum* L. exposed to salinity stress. Rhizosphere 26:100684. doi: 10.1016/j.rhisph.2023.100684

Su, Z., Ma, X., Guo, H., Sukiran, N. L., Guo, B., Assmann, S. M., et al. (2013). Flower development under drought stress: morphological and transcriptomic analyses reveal acute responses and long-term acclimation in *Arabidopsis*. Plant Cell 25, 3785–3807. doi: 10.1105/tpc.113.115428

Tardieu, F., and Simonneau, T. (1998). Variability among species of stomatal control under fluctuating soil water status and evaporative demand: modelling isohydric and anisohydric behaviours. J. Exp. Bot. 49, 419–432. doi: 10.1093/jxb/49.Special_Issue.419

Teixeira, A., Iannetta, P., Binnie, K., Valentine, T. A., and Toorop, P. (2020). Myxospermous seed-mucilage quantity correlates with environmental gradients indicative of water-deficit stress: *Plantago* species as a model. Plant Soil 446, 343–356. doi: 10.1007/s11104-019-04335-z

Tsai, A. Y. L., McGee, R., Dean, G. H., Haughn, G. W., and Sawa, S. (2021). Seed mucilage: biological functions and potential applications in biotechnology. Plant Cell Physiol. 62, 1847–1857. doi: 10.1093/pcp/pcab099

Turc, O., and Tardieu, F. (2018). Drought affects abortion of reproductive organs by exacerbating developmentally driven processes via expansive growth and hydraulics. J. Exp. Bot. 69, 3245–3254. doi: 10.1093/jxb/ery078

Vayssières, A., Boutet, S., Totozafy, J. C., Tellier, F., Winkler, M., Barreda, L., et al. (2026). Distinct metabolic signatures associated with drought response, shoot architecture, and flowering time in camelina. Plant Cell Physiol. 67, 186–201. doi: 10.1093/pcp/pcaf150

Voiniciuc, C. (2016). Quantification of the mucilage detachment from *Arabidopsis* seeds. Bio Protoc. 6:e1802. doi: 10.21769/BioProtoc.1802

Voiniciuc, C., Zimmermann, E., Schmidt, M. H. W., Günl, M., Fu, L., North, H. M., et al. (2016). Extensive natural variation in *Arabidopsis* seed mucilage structure. Front. Plant Sci. 7:803. doi: 10.3389/fpls.2016.00803

Waraich, E. A., Ahmed, Z., Ahmad, R., Saifullah, R., Shahbaz, M., and Ehsanullah (2017). Modulation in growth, development, and yield of *Camelina sativa* by nitrogen application under water stress conditions. J. Plant Nutr. 40, 726–735. doi: 10.1080/01904167.2016.1262404

Wickham, H., Averick, M., Bryan, J., Chang, W., McGowan, L., François, R., et al. (2019). Welcome to the tidyverse. J. Open Source Softw. 4:1686. doi: 10.21105/joss.01686

Yang, B., Hofmann, F., Usadel, B., and Voiniciuc, C. (2021). Seed hemicelluloses tailor mucilage properties and salt tolerance. New Phytol. 229, 1946–1954. doi: 10.1111/nph.17056

Yang, X., Baskin, C. C., Baskin, J. M., Zhang, W., and Huang, Z. (2012). Degradation of seed mucilage by soil microflora promotes early seedling growth of a desert sand dune plant. Plant Cell Environ. 35, 872–883. doi: 10.1111/j.1365-3040.2011.02459.x

Zanetti, F., Peroni, P., Pagani, E., von Cossel, M., Greiner, B. E., Krzyżaniak, M., et al. (2024). The opportunities and potential of camelina in marginal land in Europe. Ind. Crops Prod. 211:118224. doi: 10.1016/j.indcrop.2024.118224

