## Supplementary_Material for "Drought responses reshape growth, reproduction and seed traits in *Camelina sativa*"

### Supplementary Tables

**Table S1.** Statistical tests of the models reported in the Results. For each trait, the tests of accession, treatment and their interaction are shown, followed by other terms where the model includes them. F tests are Type III with sum-to-zero contrasts, with Satterthwaite denominator degrees of freedom for linear mixed models, and are given as F (numerator df, denominator df); $\chi^2$ tests are Wald tests, except the likelihood-ratio tests of A × T within each germination condition. LM, linear model; LMM, linear mixed model with plant as random intercept (and stem within plant for stem class); GLMM, generalized linear mixed model. FDR q values are adjusted across the 14 monosaccharide × fraction combinations. SSMIL, secondary stem main inflorescence length.

| Trait (model) | Accession (A) | Treatment (T) | A×T | Other terms |
|---|---|---|---|---|
| **Shoot growth and leaf physiology: $\chi^2$ (df); P for shoot area, F (df); P otherwise** | | | | |
| Projected shoot area (GLMM, AR(1) over weeks) | 17.53 (5); 0.004 | 77.86 (1); < 0.001 | 11.86 (5); 0.037 | Week (W): 448.93 (5); < 0.001<br>A×W: 70.89 (25); < 0.001<br>T×W: 189.45 (5); < 0.001<br>A×T×W: 66.05 (25); < 0.001 |
| Leaf temperature (LMM) | 4.12 (5, 23.1); 0.008 | 261.17 (1, 23.1); < 0.001 | 2.37 (5, 23.1); 0.071 | |
| Stomatal conductance (LMM, variances per treatment) | 1.28 (5, 24); 0.31 | 77.03 (1, 24); < 0.001 | 1.15 (5, 24); 0.36 | |
| **Plant architecture, seed yield and seed traits: F (df); P** | | | | |
| Secondary stem length (LMM) | 5.25 (5, 21.9); 0.003 | 92.84 (1, 22.3); < 0.001 | 1.61 (5, 21.9); 0.20 | |
| SSMIL (LMM) | 0.76 (5, 25.1); 0.58 | 286.90 (1, 25.2); < 0.001 | 1.74 (5, 25.1); 0.16 | |
| Internode distance (LMM) | 6.61 (5, 24.0); < 0.001 | 106.77 (1, 24.2); < 0.001 | 1.94 (5, 24.0); 0.12 | |
| Secondary stem density (LM) | 10.57 (5, 24); < 0.001 | 125.65 (1, 24); < 0.001 | 1.33 (5, 24); 0.28 | |
| Number of secondary stems (LM) | 43.00 (5, 24); < 0.001 | 7.51 (1, 24); 0.011 | 0.47 (5, 24); 0.79 | |
| Main stem length (LM) | 2.62 (5, 24); 0.050 | 207.24 (1, 24); < 0.001 | 1.49 (5, 24); 0.23 | |
| Basal-to-mid stem diameter ratio (LM) | 7.93 (5, 24); < 0.001 | 0.22 (1, 24); 0.64 | 1.67 (5, 24); 0.18 | |
| Seed yield per plant (LM) | 2.71 (5, 24); 0.044 | 53.44 (1, 24); < 0.001 | 2.19 (5, 24); 0.088 | |
| Seed number per plant (LM) | 2.11 (5, 24); 0.100 | 91.80 (1, 24); < 0.001 | 1.57 (5, 24); 0.21 | |
| Seed weight (LM) | 12.89 (5, 24); < 0.001 | 113.37 (1, 24); < 0.001 | 0.17 (5, 24); 0.97 | |
| Seed size (LMM) | 32.58 (5, 24.0); < 0.001 | 166.70 (1, 24.0); < 0.001 | 0.50 (5, 24.0); 0.77 | |
| **Seed mucilage: F (df); P** | | | | |
| MAS (LMM) | 8.40 (5, 24.0); < 0.001 | 1.54 (1, 24.0); 0.23 | 2.26 (5, 24.0); 0.081 | |
| MAS : seed size ratio (LMM) | 8.10 (5, 24.0); < 0.001 | 41.68 (1, 24.0); < 0.001 | 2.08 (5, 24.0); 0.10 | |
| MAS by stem class (LMM) | 8.35 (5, 24.0); < 0.001 | 2.05 (1, 24.0); 0.17 | 1.91 (5, 24.0); 0.13 | Stem class (S): 0.41 (2, 46.8); 0.67<br>A×S: 2.06 (10, 46.8); 0.048<br>T×S: 0.46 (2, 46.8); 0.63<br>A×T×S: 0.57 (10, 46.8); 0.83 |
| MAS : seed size ratio by stem class (LMM) | 7.36 (5, 24.0); < 0.001 | 40.36 (1, 24.0); < 0.001 | 1.67 (5, 24.0); 0.18 | Stem class (S): 2.46 (2, 46.8); 0.096 |

| Trait (model) | Accession (A) | Treatment (T) | A×T | Other terms |
|---|---|---|---|---|
| | | | | A×S: 2.83 (10, 46.8); 0.008<br>T×S: 1.60 (2, 46.8); 0.21<br>A×T×S: 0.58 (10, 46.8); 0.82 |
| **Monosaccharide composition (LM per monosaccharide and fraction, plant means): F (df); P; FDR q for treatment** | | | | |
| Rha, non-adherent | 18.60 (5, 22); < 0.001 | 12.35 (1, 22); 0.002 | 8.77 (5, 22); < 0.001 | q (treatment) = 0.021 |
| Ara, non-adherent | 19.19 (5, 22); < 0.001 | 0.96 (1, 22); 0.34 | 2.60 (5, 22); 0.054 | q (treatment) = 0.55 |
| Gal, non-adherent | 18.03 (5, 22); < 0.001 | 3.73 (1, 22); 0.067 | 4.82 (5, 22); 0.004 | q (treatment) = 0.13 |
| Glc, non-adherent | 1.61 (5, 22); 0.20 | 5.89 (1, 22); 0.024 | 1.02 (5, 22); 0.43 | q (treatment) = 0.067 |
| Xyl, non-adherent | 2.12 (5, 22); 0.10 | 0.24 (1, 22); 0.63 | 1.08 (5, 22); 0.40 | q (treatment) = 0.84 |
| Man, non-adherent | 1.50 (5, 22); 0.23 | 9.41 (1, 22); 0.006 | 0.27 (5, 22); 0.92 | q (treatment) = 0.026 |
| GalA, non-adherent | 4.76 (5, 22); 0.004 | 0.89 (1, 22); 0.36 | 0.40 (5, 22); 0.84 | q (treatment) = 0.55 |
| Rha, adherent | 1.00 (5, 23); 0.44 | 0.18 (1, 23); 0.68 | 0.18 (5, 23); 0.97 | q (treatment) = 0.84 |
| Ara, adherent | 0.96 (5, 23); 0.46 | 0.02 (1, 23); 0.90 | 0.47 (5, 23); 0.79 | q (treatment) = 0.90 |
| Gal, adherent | 2.86 (5, 23); 0.038 | 4.22 (1, 23); 0.051 | 1.04 (5, 23); 0.42 | q (treatment) = 0.12 |
| Glc, adherent | 4.41 (5, 23); 0.006 | 11.02 (1, 23); 0.003 | 4.31 (5, 23); 0.007 | q (treatment) = 0.021 |
| Xyl, adherent | 1.53 (5, 23); 0.22 | 0.11 (1, 23); 0.74 | 1.29 (5, 23); 0.30 | q (treatment) = 0.84 |
| Man, adherent | 1.92 (5, 22); 0.13 | 0.08 (1, 22); 0.78 | 0.62 (5, 22); 0.69 | q (treatment) = 0.84 |
| GalA, adherent | 1.73 (5, 23); 0.17 | 7.31 (1, 23); 0.013 | 0.97 (5, 23); 0.45 | q (treatment) = 0.044 |
| **Fraction difference (non-adherent – adherent per plant, LM): F (df); P** | **Fraction** | **T × fraction** | | **FDR q** |
| Rha | 353.94 (1, 22); < 0.001 | 2.09 (1, 22); 0.16 | | < 0.001 |
| Ara | 23.25 (1, 22); < 0.001 | 0.02 (1, 22); 0.90 | | < 0.001 |
| Gal | 183.69 (1, 22); < 0.001 | 8.05 (1, 22); 0.010 | | < 0.001 |
| Glc | 28.21 (1, 22); < 0.001 | 9.62 (1, 22); 0.005 | | < 0.001 |
| Xyl | 97.23 (1, 22); < 0.001 | 0.15 (1, 22); 0.70 | | < 0.001 |
| Man | 20.16 (1, 21); < 0.001 | 9.14 (1, 21); 0.006 | | < 0.001 |
| GalA | 0.68 (1, 22); 0.42 | 3.78 (1, 22); 0.065 | | 0.42 |
| **PERMANOVA on clr-transformed composition: pseudo-F (df); $R^2$; P** | **Accession** | **Treatment** | **A×T** | |
| Non-adherent mucilage | 6.50 (5, 22); $R^2$ = 0.44; < 0.001 | 10.49 (1, 22); $R^2$ = 0.14; < 0.001 | 1.89 (5, 22); $R^2$ = 0.13; 0.0487 | |
| Adherent mucilage | 3.18 (5, 22); $R^2$ = 0.31; 0.0022 | 3.14 (1, 22); $R^2$ = 0.06; 0.0455 | 2.08 (5, 22); $R^2$ = 0.20; 0.0405 | |
| **Germination (binomial GLMM, Wald $\chi^2$ (df); P)** | | | | |
| Final germination, day 3 (NaCl −0.6, NaCl −1.2, PEG −0.6 MPa) | 14.31 (5); 0.014 | 22.40 (1); < 0.001 | 10.18 (5); 0.070 | Condition (C): 693.06 (2); < 0.001<br>A×C: 12.83 (10); 0.23<br>T×C: 0.96 (2); 0.62 |
| Germination speed, day 1 (water, NaCl −0.6, NaCl −1.2, PEG −0.4 MPa) | 14.88 (5); 0.011 | 4.22 (1); 0.040 | 5.96 (5); 0.31 | Condition (C): 389.33 (3); < 0.001<br>A×C: 61.07 (15); < 0.001<br>T×C: 4.94 (3); 0.18 |
| **A×T within each condition, day 3 (likelihood-ratio test, $\chi^2$ (df); P)** | | | **A×T** | |
| NaCl −0.6 MPa | | | 3.2 (5); 0.67 | |
| NaCl −1.2 MPa | | | 9.9 (5); 0.079 | |
| PEG −0.6 MPa | | | 16.4 (5); 0.006 | |

**Table S2.** Number of secondary stems analyzed with main stem length as a covariate, and mucilage area per seed (MAS) analyzed with seed size as a covariate (analyses of covariance). Both models include accession, treatment and their interaction; the MAS model is a linear mixed model with plant as random intercept, fitted by REML, with Satterthwaite degrees of freedom. Covariates were centred on their mean. Drought effects are the % change relative to control plants, at the mean main stem length (51.5 cm) or the mean log seed size, with 95% CI (delta method); P values per accession are Holm-adjusted across the six accessions. Seed size is the imbibed seed area used for the MAS : seed size ratio.

| | Number of secondary stems with main stem length as covariate (LM, 36 plants) | log MAS with log seed size as covariate (LMM, 5,572 seeds from 36 plants)[a] |
|---|---|---|
| Type III F tests: F (df); P | | |
| Covariate | 0.52 (1, 23); 0.48 | 3017 (1, 5468.2); < 0.001 |
| Accession | 40.71 (5, 23); < 0.001 | 6.74 (5, 24.1); < 0.001 |
| Treatment | 0.04 (1, 23); 0.85 | 88.80 (1, 24.7); < 0.001 |
| Accession × treatment | 0.43 (5, 23); 0.83 | 1.48 (5, 23.9); 0.23 |
| Covariate × treatment[b] | P = 0.27 | P = 0.035[b] |
| Covariate slope (95% CI) | | |
| Slope | −0.07 stems per cm (−0.28 to 0.13) | 2.24 (2.16 to 2.32)[c] |
| Drought effect, % (95% CI); P | | |
| Without covariate | +4.5 (+1.1 to +8.0); 0.011 | −13.8 (−30.4 to +6.8); 0.17[d] |
| At equal main stem length or seed size[e] | +1.0 (−9.0 to +12.1); 0.85 | −47.3 (−54.2 to −39.4); < 0.001 |
| Drought effect at equal main stem length or seed size, by accession, % (95% CI); P (Holm) | | |
| CAM22 | +3.0 (−8.0 to +15.3); 1.00 | −60.1 (−71.6 to −43.9); < 0.001 |
| CAM70 | −2.0 (−16.0 to +14.2); 1.00 | −41.0 (−58.1 to −17.0); 0.012 |
| CAM131 | −2.0 (−14.7 to +12.7); 1.00 | −36.6 (−54.9 to −10.8); 0.012 |
| CAM190 | −0.2 (−11.2 to +12.3); 1.00 | −42.1 (−58.8 to −18.6); 0.012 |
| CAM214 | +4.3 (−6.7 to +16.6); 1.00 | −40.5 (−57.7 to −16.4); 0.012 |
| CAM230 | +1.5 (−12.5 to +17.7); 1.00 | −58.3 (−70.4 to −41.3); < 0.001 |

[a] 52 of 5,624 seeds (0.9%; 31 control, 21 drought) with zero or negative MAS were excluded, as in all mucilage analyses.

[b] Tested by adding the covariate × treatment interaction. For MAS, the slope was slightly steeper in drought than in control seeds (2.33 vs 2.15), but the drought effect at the mean seed size was the same in this model (−47.3%, 95% CI −54.2 to −39.3%). In an ANCOVA on plant means (36 plants), the slope did not differ between treatments (P = 0.79) and the drought effect was −45.4% (−56.2 to −32.0%).

[c] The slope is above 1 (P < 0.001), both among seeds within a plant (2.25) and among plants (2.08, 95% CI 1.32 to 2.84).

[d] Geometric mean (log scale).

[e] Drought and control plants did not overlap in main stem length (38.0–49.8 cm vs 50.0–67.5 cm), so the adjusted effect on the number of secondary stems relies on extending the common slope beyond the data.

**Table S3. Associations of seed traits and mucilage with progeny germination (final germination, day 3), and of mucilage composition with seed traits.** Among plants: binomial GLMMs fitted separately for each condition, with accession and treatment as fixed effects, an observation-level random intercept and the standardized predictor; MAS was fitted together with imbibed seed size. Cells give the odds ratio per standard deviation (95% CI); likelihood-ratio $\chi^2$ (1); P. Among accessions: Spearman correlation between the drought change in germination and the drought change in the predictor (six accessions); cells give ρ; P. FDR-adjusted P values are Benjamini–Hochberg adjusted within each group of tests (seed traits, 6 tests; monosaccharides, 42 tests; composition vs seed traits, 28 tests per level). The Glc association under NaCl −1.2 MPa rests mainly on CAM230 (without it: OR 0.70, P = 0.12, 29 plants). AM, adherent mucilage; NAM, non-adherent mucilage; Glc, glucose; MAS, mucilage area per seed.

| Predictor | NaCl −0.6 MPa | NaCl −1.2 MPa | PEG −0.6 MPa | FDR-adjusted P |
|---|---|---|---|---|
| **Among plants within accessions and treatments: OR (95% CI); χ² (1); P** | | | | |
| Seed size | 0.76 (0.29–2.01); 0.31; 0.58 | 0.21 (0.05–0.84); 4.71; 0.030 | 0.81 (0.39–1.68); 0.33; 0.57 | ≥ 0.18 |
| Seed weight | 0.98 (0.49–1.98); 0.00; 0.96 | 0.61 (0.21–1.82); 0.77; 0.38 | 0.72 (0.40–1.30); 1.19; 0.28 | ≥ 0.70 |
| MAS (with seed size as covariate) | 2.08 (0.96–4.51); 3.55; 0.060 | 3.03 (1.35–6.80); 6.80; 0.009 | 1.02 (0.66–1.58); 0.01; 0.93 | 0.027 (NaCl −1.2 MPa); ≥ 0.090 otherwise |
| MAS × treatment: χ² (1); P | 0.05; 0.82 | 0.52; 0.47 | 0.36; 0.55 | |
| Glc, AM | 1.11 (0.77–1.60); 0.32; 0.57 | 0.42 (0.25–0.70); 11.27; < 0.001 | 0.79 (0.61–1.02); 3.13; 0.077 | 0.033 (NaCl −1.2 MPa); ≥ 0.40 otherwise |
| Other monosaccharides (6 in AM, 7 in NAM) | | | | ≥ 0.31 |
| **Among accessions, drought change: Spearman ρ; P** | | | | |
| Seed size | 0.53; 0.28 | 0.03; 1.00 | 0.71; 0.14 | ≥ 0.38 |
| Seed weight | 0.62; 0.19 | 0.49; 0.36 | 0.71; 0.14 | ≥ 0.38 |
| MAS | −0.44; 0.38 | 0.26; 0.66 | 0.03; 1.00 | ≥ 0.99 |
| Monosaccharides (7 in AM, 7 in NAM) | | | | ≥ 0.83 |
| **Mucilage composition (7 monosaccharides in AM and NAM) vs seed size and seed weight** | | | | |
| Among plants (LM with accession and treatment) | | | | ≥ 0.16 |
| Among accessions (Spearman, drought change) | | | | ≥ 0.08 |

### Supplementary Figures

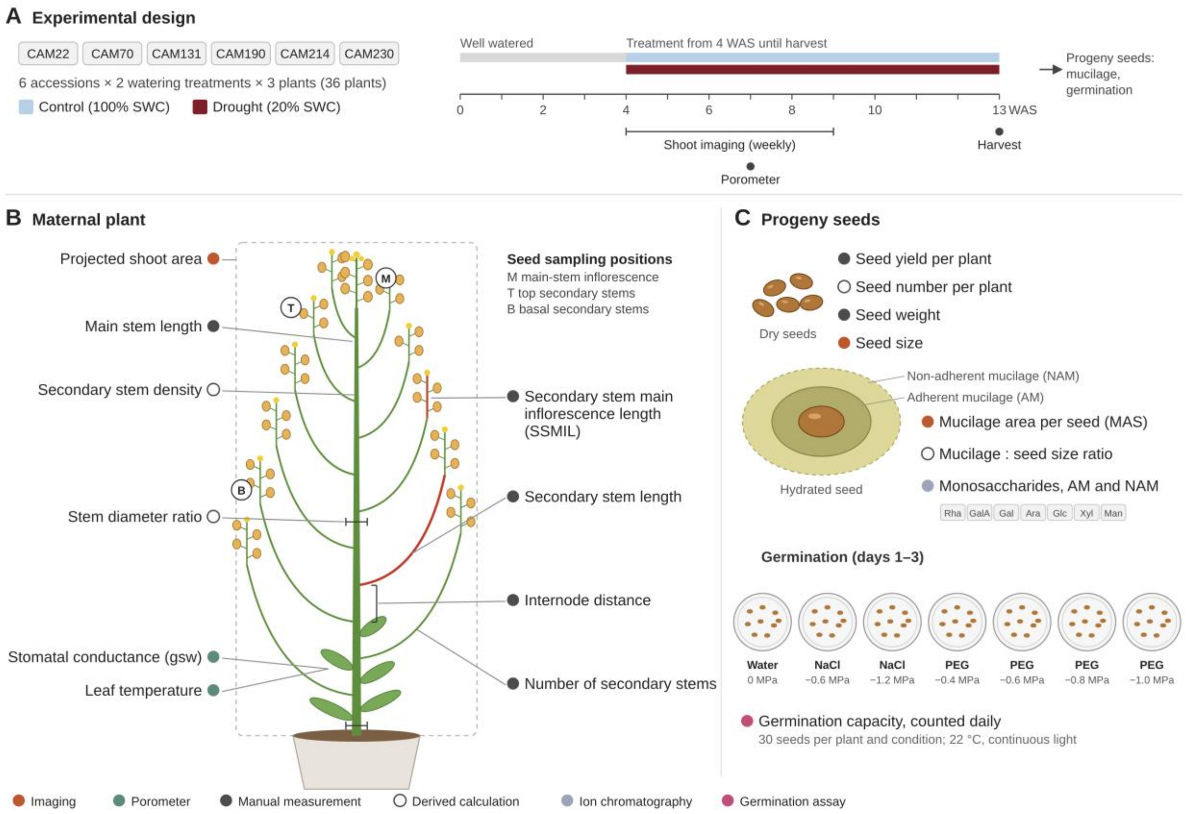


**Figure S1. Experimental design and traits measured on maternal *Camelina* plants and their progeny seeds. (A)** Six accessions were grown with three plants per accession and treatment (36 plants). From 4 weeks after sowing (WAS) until harvest at 13 WAS, plants were kept at 100% (control) or 20% (drought) soil water content (SWC). Shoots were imaged weekly, and stomatal conductance and leaf temperature were measured once with a porometer. **(B)** Traits measured on the maternal plant. Red lines mark the stem lengths measured as secondary stem length and secondary stem main inflorescence length (SSMIL); M, T and B mark the seed sampling positions (main stem main inflorescence, top and basal secondary stems). **(C)** Traits measured on the progeny seeds: seed yield and seed number per plant, seed weight and size, mucilage area of the hydrated seed, the monosaccharide composition of adherent (AM) and non-adherent (NAM) mucilage, and germination in water, NaCl and PEG at the water potentials shown (30 seeds per plant and condition, 22 °C, continuous light, scored daily for 3 days). Dot colors show how each trait was measured; open circles are traits calculated from other measurements.

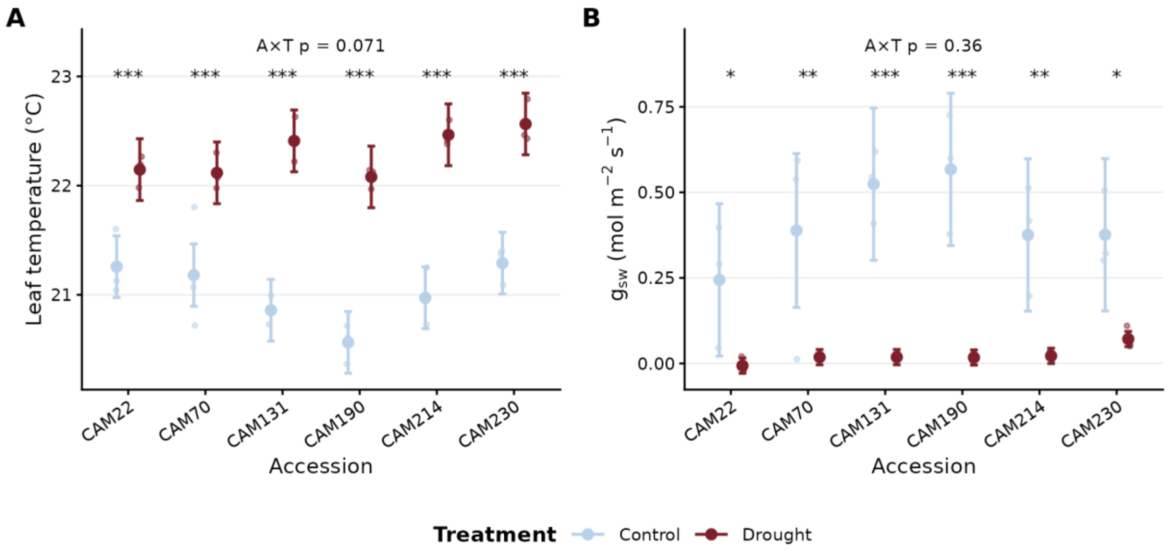


**Figure S2. Leaf temperature and stomatal conductance of each *Camelina* accession under control and drought conditions. (A)** Leaf temperature and **(B)** stomatal conductance (gsw) per accession, from the same models as Figure 1B–C (estimated marginal means with 95% CI; small dots are plant means). Stars give the Holm-adjusted drought vs control contrast within each accession (* $P < 0.05$, ** $P < 0.01$, *** $P < 0.001$); the subtitle gives the accession × treatment test.

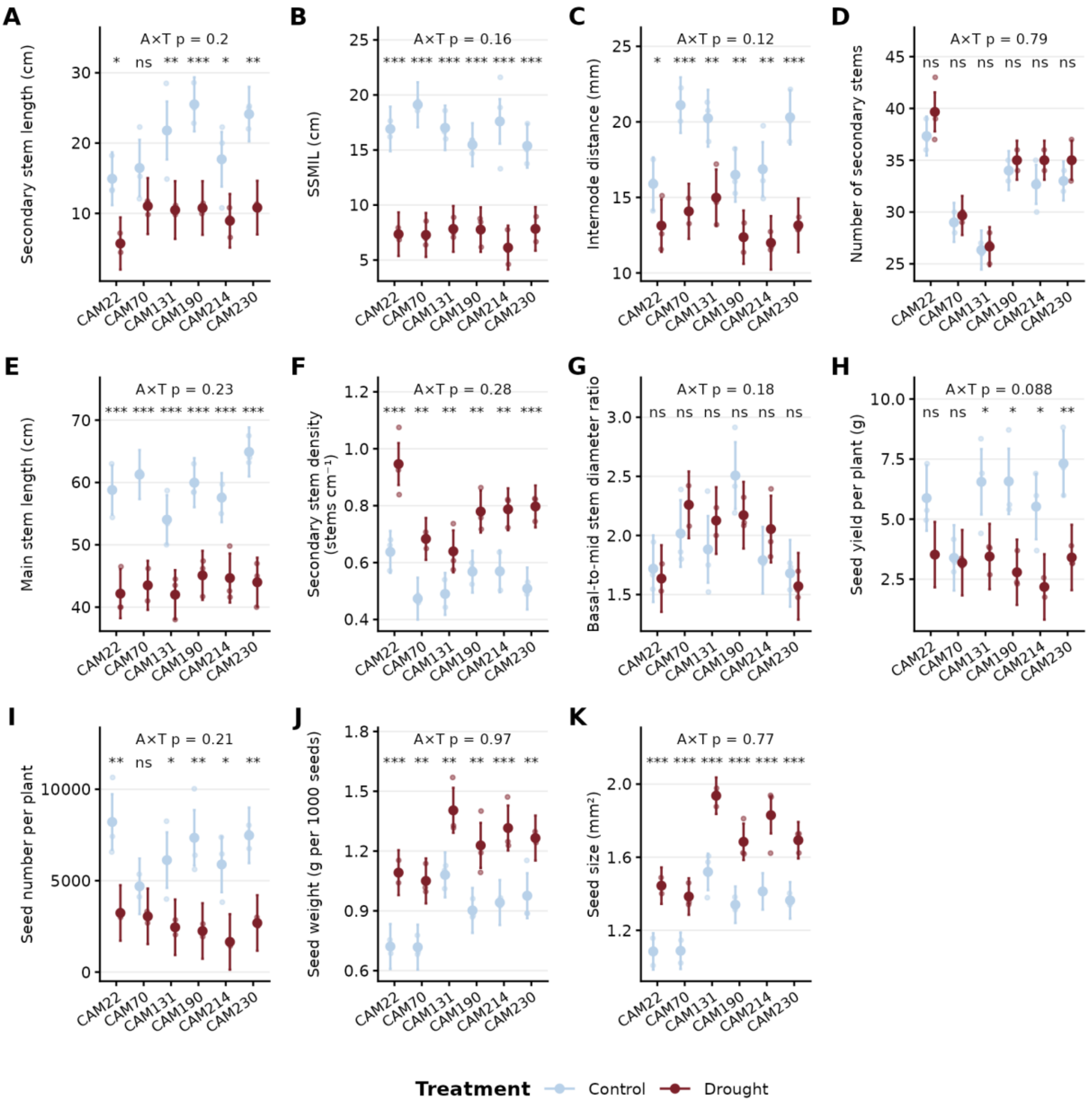


**Figure S3. Maternal plant architecture, seed yield and progeny seed traits of each *Camelina* accession under control and drought conditions. (A)** Secondary stem length, **(B)** secondary stem main inflorescence length (SSMIL), **(C)** internode distance, **(D)** number of secondary stems, **(E)** main stem length, **(F)** secondary stem density, **(G)** basal-to-mid stem diameter ratio, **(H)** seed yield per plant, **(I)** seed number per plant, **(J)** seed weight and **(K)** seed size, from the same models as Figure 2 (estimated marginal means with 95% CI; small dots are plant means). Stars give the Holm-adjusted drought vs control contrast within each accession (* $P < 0.05$, ** $P < 0.01$, *** $P < 0.001$; ns, not significant). A × T, accession × treatment interaction.

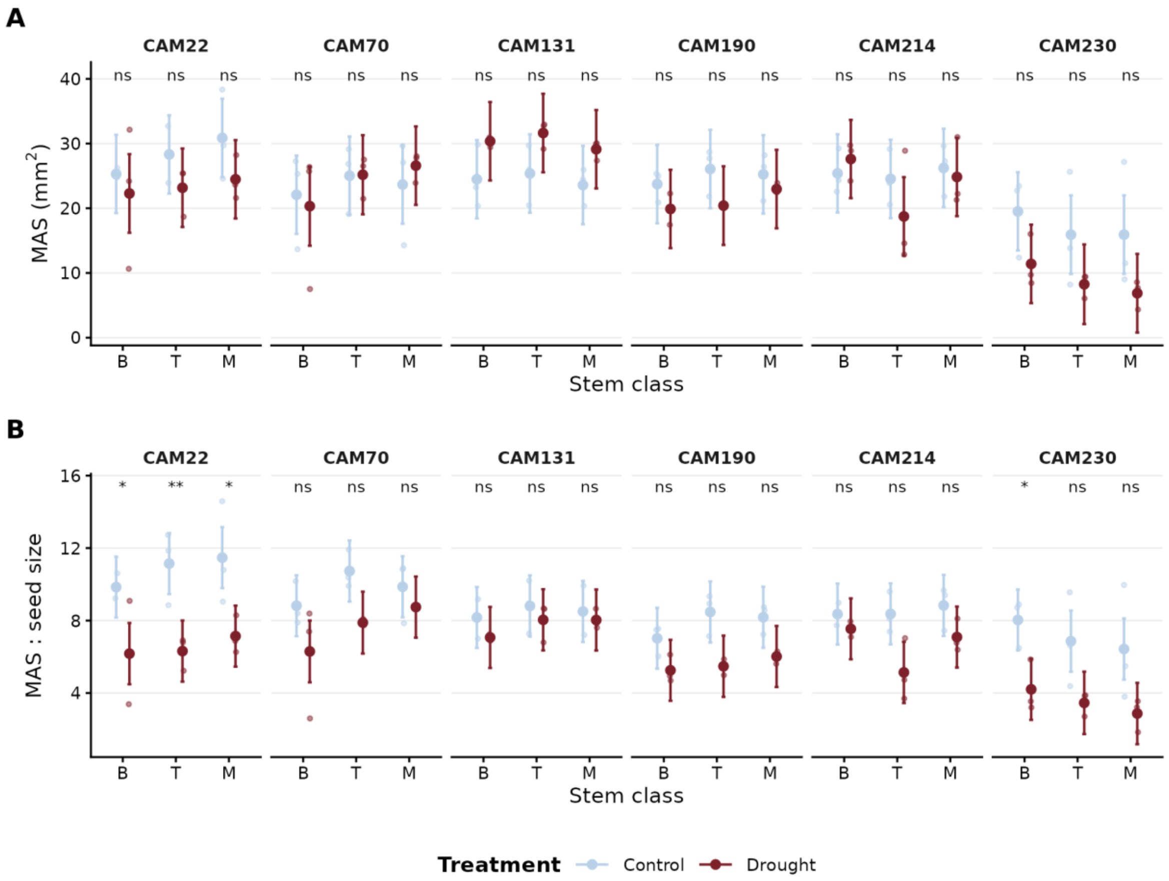


**Figure S4. Seed mucilage by stem class in each *Camelina* accession under control and drought conditions. (A)** Mucilage area per seed (MAS) and **(B)** MAS : seed size ratio by stem class (B, basal secondary stem; T, top secondary stem; M, main stem main inflorescence) for each accession, from the same joint models as Figure 3C–D (estimated marginal means with 95% CI; small dots are stem means). Stars give the drought vs control contrast within each accession and stem class, Holm-adjusted across all 18 contrasts per trait (* P < 0.05, ** P < 0.01; ns, not significant).

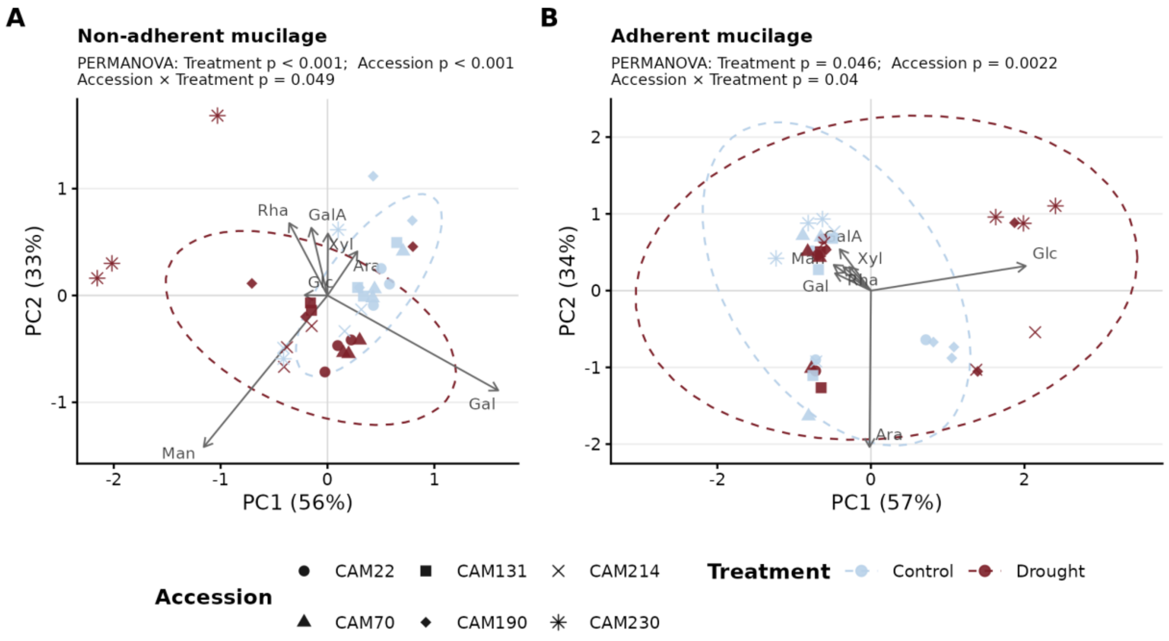


**Figure S5. Principal component analysis of the monosaccharide composition of (A) non-adherent and (B) adherent seed mucilage from control and drought-grown plants of six *Camelina* accessions.** Each point is one plant (mean of up to three capsule subsamples); color shows the maternal treatment and symbol the accession. Molar percentages of Rha, Ara, Gal, Glc, Xyl, Man and GalA were centred log-ratio transformed before analysis; arrows show the loading of each sugar and dashed ellipses the 95% normal-theory region per treatment. P values are from PERMANOVA on Aitchison distance (Euclidean distance of clr data; 9,999 permutations; sequential terms accession, treatment, accession × treatment). Multivariate dispersion did not differ between treatments (betadisper, non-adherent P = 0.12, adherent P = 0.14).

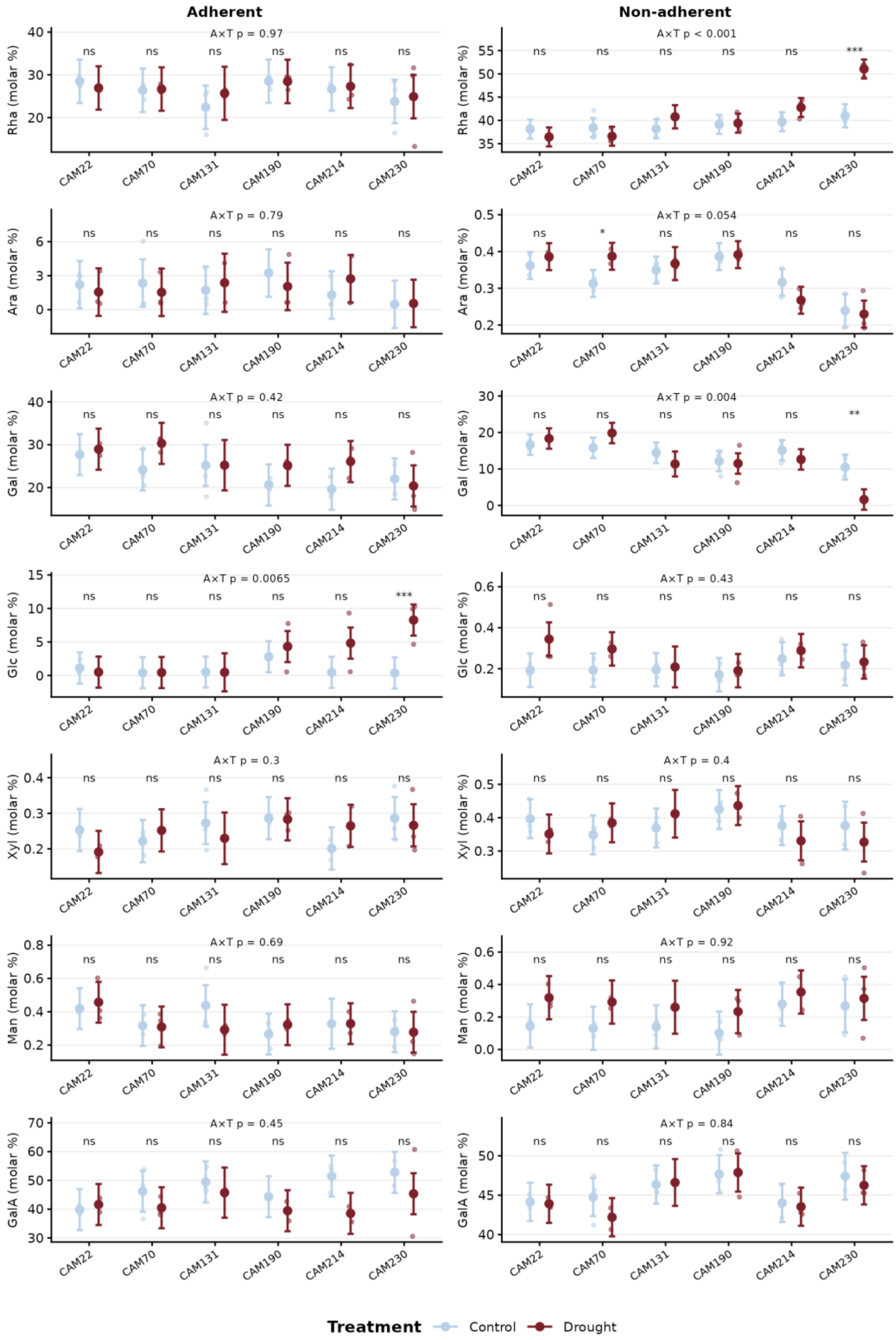


**Figure S6. Monosaccharide composition of adherent and non-adherent mucilage in each *Camelina* accession under control and drought conditions.** Molar percentage of each sugar per accession, from the same models as Figure 4 (estimated marginal means with 95% CI). Small dots are plant means and stars give the Holm-adjusted drought vs control contrast within each accession; A × T, accession × treatment test.
